# Stretch-evoked motor responses in the brainstem are modulated by task instructions

**DOI:** 10.64898/2026.04.03.716388

**Authors:** Rebecca C. Nikonowicz, Neha A. Reddy, Michelle C. Medina, Molly G. Bright, Fabrizio Sergi

**Affiliations:** Department of Biomedical Engineering, UD College of Engineering, University of Delaware, Newark, DE 19716, USA; Department of Physical Therapy and Human Movement Sciences, Feinberg School of Medicine, Northwestern University, Chicago, IL 60611, USA; Department of Biomedical Engineering, McCormick School of Engineering and Applied Sciences, Northwestern University, Evanston, IL 60208, USA; Department of Mechanical Engineering, UD College of Engineering, University of Delaware, Newark, DE, 19713, USA

## Abstract

Reliable noninvasive measurement of human brainstem activity during motor control remains challenging due to small anatomical structures and physiological noise, yet it is essential for understanding descending contributions to rapid feedback control. We used brainstem-optimized whole brain functional magnetic resonance imaging (fMRI) to examine whether task-dependent modulation of stretch-evoked motor responses in humans are associated with changes in activation within reticulospinal regions of the human brainstem. A behavioral validation experiment (N=10), in which participants were instructed to resist or yield to brief wrist perturbations, confirmed task-dependent modulation of long-latency responses (LLRs). In a separate imaging cohort (N=26), participants performed the same tasks during fMRI using an MRI-compatible robotic device and a multi-echo acquisition with physiological noise compensation. Imaging analyses revealed greater blood-oxygen-level-dependent signal during Resist compared to Yield while controlling for background contraction and proprioceptive input, with activation distributed bilaterally across the pons and medulla in regions consistent with major reticulospinal nuclei. Laterality analyses demonstrated a rostrocaudal gradient, with relatively ipsilateral-biased activation in the medulla shifting toward more contralateral patterns in the pons. These findings indicate that instruction-dependent modulation of stretch-evoked motor responses is associated with measurable changes in human brainstem activation, providing evidence that reticulospinal regions contribute to task-dependent feedback control.

## 1 Introduction

The reticulospinal tract (RST) is a descending motor pathway that plays a critical role in postural control, gross limb movements, and motor recovery following neurological injury. Unlike the corticospinal tract (CST), which enables fine, distal control, the RST contributes to broader, bilateral muscle activation and appears to compensate for corticospinal damage in individuals with stroke or spinal cord injury (McPherson et al., 2018; Baker, 2011). Despite its functional significance, the RST remains poorly characterized in humans, largely due to its deep location in the brainstem and the absence of reliable, non-invasive techniques to quantify its activity. As a result, direct evidence of RST engagement during voluntary or task-modulated behavior in humans has been elusive.

One promising strategy to probe reticulospinal activity involves the study of long-latency responses (LLRs), stereotypical muscle responses occurring 50-100 ms after a muscle-stretching perturbation and likely involving both spinal and supraspinal processing (Pruszynski et al., 2011; Kurtzer, 2014). LLRs are modulated by task goals and instructions. Specifically, LLR amplitude (LLRa) will be higher if a participant attempts to resist a perturbation as opposed to yield to it (Lewis et al., 2006). This task-dependent gain scaling likely reflects supraspinal engagement and in fact, many studies have concluded that LLRs are generated and/or modulated by the primary motor cortex (M1), sensorimotor cortex (S1), premotor cortex, somatosensory areas, and secondary somatosensory cortex (Spieser et al., 2013; C. Capaday, 1991; Kourtis et al., 2008; Palmer and Ashby, 1992; Soteropoulos and Baker, 2020; Marsden et al., 1977).

While these studies emphasize the cortical contributions to LLRs, Shemmel et al. showed that transcranial magnetic stimulation (TMS) suppression of M1 does not diminish LLRa during a Resist condition (Shemmell et al., 2009), suggesting the goal-dependent component of LLRs is modulated elsewhere in the brain. One plausible candidate for processing such goal-dependent component of the LLRs is the reticular formation (RF), which originates the RST. The RF has been strongly implicated in the startle response via the “StartReact” paradigm in which a startling acoustic stimulus triggers a faster than normal preplanned motor action (Valls-Solé et al., 1999; Carlsen et al., 2011). These StartReact responses occur at latencies under 70 ms, faster than typical voluntary reaction times, and are thought to rely on subcortical circuits rather than cortical drive alone (Valls-Solé et al., 1999; Carlsen et al., 2011). Supporting this theory, individuals with stroke or degenerated CST have delayed voluntary reactions but normal onset of StartReact (Honeycutt and Perreault, 2012). Further, Ravichandran et al. demonstrated a similar pattern of activity in the LLR epoch for participants attempting to quickly initiate an elbow movement with (1) a startling acoustic stimulus and (2) a perturbation, suggesting common underlying pathways (Ravichandran et al., 2013). Taken together, these findings implicate the RF, and by extension the RST, as a likely contributor to task-dependent LLR modulation in humans, offering a non-invasive behavioral probe of RST function.

Despite growing evidence implicating the RST in rapid, goal-directed motor responses, most human studies have inferred its involvement indirectly through behavioral paradigms such as StartReact or through peripheral physiological measures including electromyography (EMG) amplitude and response latency. To the authors’ knowledge, no study has measured human brainstem activation associated with task-dependent modulation of LLRs during wrist perturbations. This gap reflects both anatomical and technical challenges: the RF lies deep within the brainstem and consists of small, densely packed nuclei, making it difficult to isolate function using noninvasive neuroimaging due to limited spatial resolution and substantial physiological noise in this region.

To overcome these challenges, our group has developed StretchfMRI, a novel paradigm combining robotic perturbations, EMG, and functional magnetic resonance imaging (fMRI) to elicit LLRs and record muscle (EMG) and neural (fMRI) activity simultaneously (Zon- nino et al., 2019). Preliminary studies have shown that neural activity associated with LLRs in the wrist flexors and extensors can be reliably measured in the RF (Zonnino et al., 2021; Nikonowicz and Sergi, 2024). Our approach leverages the Dual Motor StretchWrist (DMSW), an MRI-compatible robotic device that delivers controlled mechanical perturbations to the wrist, alongside brainstem-optimized whole-brain fMRI acquisition and analysis. By delivering rapid perturbations paired with “Yield” and “Resist” task instructions, we use LLR modulation as a behavioral probe to differentially engage descending motor pathways. Combined with region-specific hemodynamic response modeling, denoising tailored to the brainstem and a multi-echo whole-brain fMRI sequence optimized to increase signal-to-noise in the brainstem (Reddy et al., 2024a), this method enables the visualization of neural activation in the RF. Using this experimental setup, we conducted a study to test two hypotheses: (H1) stretch-evoked fMRI signal in the RF is larger when participants are instructed to “re-sist” a perturbation compared to “yield” and (H2) brainstem activations show a rostrocaudal lateralization gradient that is consistent with the double reciprocal model of brainstem motor control (Davidson et al., 2004; Davidson and Buford, 2006; Sakai et al., 2009). Derived primarily from animal studies, this model proposes that the ipsilateral reticular formation preferentially facilitates flexor activation, while the contralateral reticular formation preferentially facilitates extensor activation. Along the pons-to-medulla axis, the model further predicts a lateralization gradient with flexor activation becoming increasingly ipsilateral in medullary regions. Although the framework provides a useful basis for interpreting brainstem motor organization, the extent to which these patterns generalize to human brainstem activation remains unclear. In addition to these primary hypotheses, we report behavioral and physiological validation outcomes, including EMG measures of stretch reflex modulation and whole-brain fMRI results to contextualize the brainstem findings within broader motor network activity. As such, this study aims to provide the first direct evidence of brainstem activation modulated by fast-feedback motor task demands in humans, offering a new, non-invasive approach to studying reticulospinal contributions to motor control.

## 2 Materials & Methods

### 2.1 Participants

10 healthy individuals (age: 25 *±* 4.45, 7M/3F) and 26 healthy individuals (age: 24 *±* 3.41, 12M/14F) participated in Experiment 1 (conducted in a mock MRI scanner) and Experiment 2 (conducted in an MRI scanner), respectively. All participants self-reported as right-handed and free from neurological disorders and orthopedic/pain conditions affecting the right wrist. Written informed consent was obtained from all participants for being included in the study. The study was approved by the Investigation Review Board of the University of Delaware under protocol no. 1097082-11 and conducted in accordance with the Declaration of Helsinki.

### 2.2 Dual Motor StretchWrist

A one degree-of-freedom robot, the Dual Motor StretchWrist (DMSW), was used to deliver precise extension perturbations to the right wrist to condition LLRs in the wrist flexors. The DMSW features two ultrasonic piezoelectric motors (EN60 Motor Shinsei Motor Inc., Japan) connected in parallel and has a maximum torque of 6 Nm, range-of-motion of 110 degrees, and peak, unloaded velocity of 900 degrees/second. The capstan transmission allows for motion transfer from the motors to the end effector and the entire device is manufactured with MR-compatible materials including 3D printed plastic (RS- F2-FPWH-04, Formlabs, Inc., MA, USA), bronze and brass fasteners and structural supports, ceramic bearings (Boca Bearings, Boynton Beach, FL, USA), and a six axis, MR-compatible force-torque sensor (Mini27Ti, ATI Industrial Automation, Apex, NC). The device and its control are described in detail in Nikonowicz et al. 2024 (Nikonowicz and Sergi, 2024) (Fig. 1).

**Figure 1:**
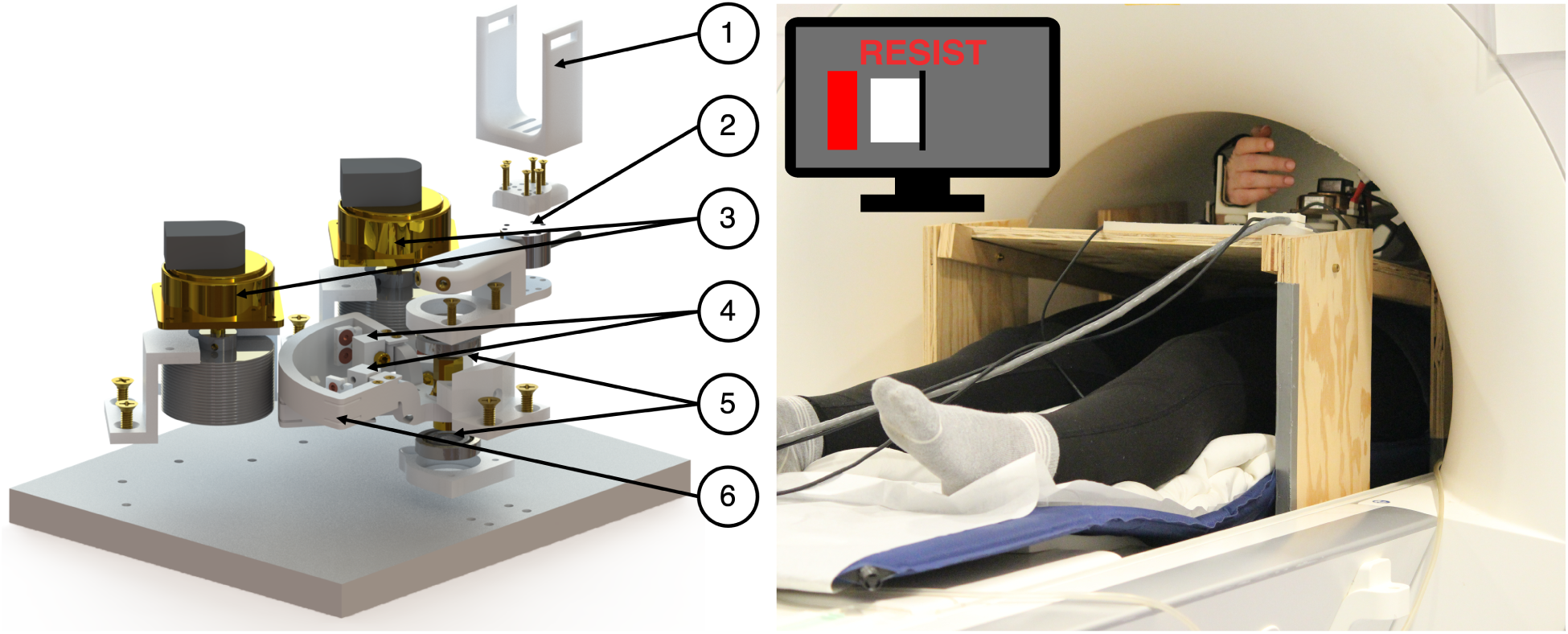
(left) Exploded view of the modeled Dual Motor StretchWrist with labels for (1) hand support, (2) MR-compatible force-torque sensor, (3) MR-compatible piezoelectric ultrasonic motors, (4) tensioning mechanisms, (5) ceramic support bearings, and (6) capstan arc. (right) A participant and the DMSW inside of the MRI scanner and a sample view of the Resist cued task.

### 2.3 EMG in the Mock MRI Scanner

Electromyography (EMG) data were collected in Experiment 1 using only surface electrodes (Delsys Trigno Duo, Natick, MA) at a sampling rate of 1926 Hz. EMG signals were recorded from the flexor carpi radialis (FCR) and extensor carpi ulnaris (ECU).

### 2.4 Yield-Resist Task

During task sessions, participants were exposed to a blocked design of perturbations to the right wrist. Before each perturbation, participants were cued via a visual interface to produce a flexion torque of 0.2*±*0.035 Nm (Fig.2(A)). After holding this torque for a duration randomly selected from a uniform distribution with extremes 400 and 800 ms, a perturbation was delivered in the wrist extension direction, resulting in stretch of the wrist flexor muscles. Perturbations were delivered at either 150 deg/s or 35 deg/s for a duration of 200 ms. After the perturbation, participants received visual feedback on their measured torque (details below) for 2 seconds before being returned to the start position at 35 deg/s. Once returned to start, the next cued flexion torque appeared immediately. Each perturbation trial lasted approximately 3.75 seconds.

**Figure 2:**
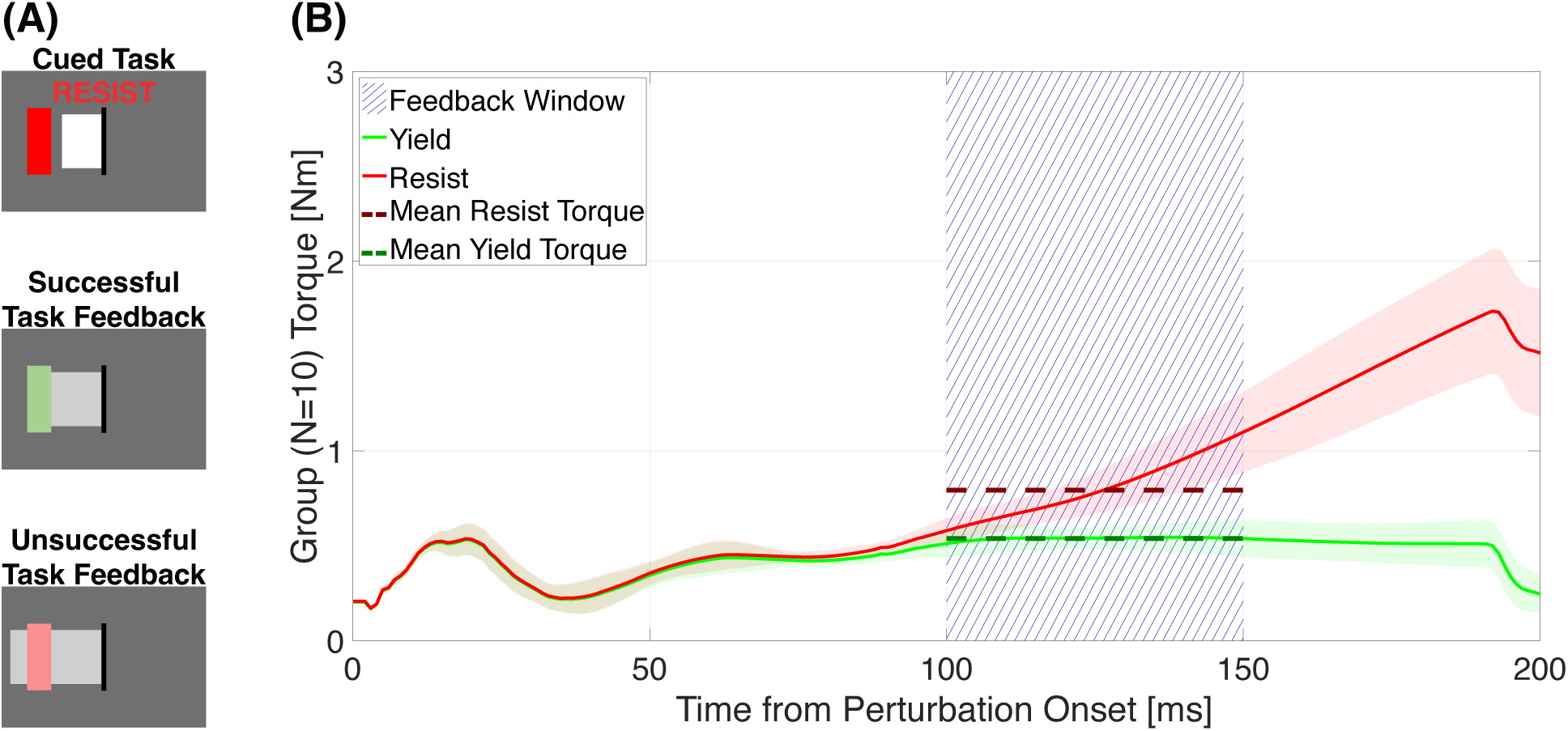
(A) Sample participant views during the task. Cued Task: what the participant sees when the task is cued, before the perturbation. The red box is the target, the white box is their live measured flexion torque, the black line is 0 Nm and the task instruction is at the top. Successful Task Feedback: what the participant sees after a perturbation when they applied the desired level of flexion torque during the perturbation where the green box is the successfully met target and the gray box is the average torque measured in the [100 150] ms post perturbation onset. Unsuccessful Task Feedback: what the participant sees after a perturbation when they applied too much torque during the perturbation where the pink box is the unmet target. (B) Average of the torque measured by the DMSW during perturbations in Experiment 1, averaged across perturbations and participants. Thick lines represent the average and shaded areas represent *±*1 s.d. indicating within-group variance. The area where torque was averaged and then displayed to participants is highlighted by blue hatch marks.

#### Task Instructions

In both experiments, participants were exposed to 3 perturbation conditions: Yield, Resist, and Slow. During Yield and Resist conditions, participants were instructed to either “Yield” to the perturbation or “Resist” the perturbation and perturbations were delivered at 150 deg/s. During Slow conditions, participants were instructed to “Yield” to the perturbation and perturbations were delivered at 35 deg/s. This Slow condition was included as it is known to minimally/not elicit stretch responses in wrist muscles (Weinman et al., 2021; Nikonowicz and Sergi, 2024).

#### Task Length

In Experiment 1, participants completed one task session. In Experiment 2, participants completed two task sessions. Each session comprised 5 blocks of each condition (15 total blocks). Each block consisted of 8 perturbations of the same condition lasting approximately 30 s, resulting in a total of 40 perturbations per condition, or a total of 120 total perturbations in each task session. Block order was randomized ([Y R R S Y Y R S Y R S S Y R S]) but held consistent across sessions and participants. In Experiment 2, each block was interleaved with 30 s rest periods. For each participant, the total number of volumes collected varied, but on average, each session lasted approximately 16 minutes. A Familiarization session preceded the task, during which participants experienced 40 perturbations per condition to become comfortable with the robotic interface and task instructions.

#### Task Torque Feedback

During the task sessions, visual feedback was provided after all perturbations to standardize resistance level in a subject-specific way. Visual feedback was provided based on the average torque value measured in the time comprised between 100 and 150 ms post-perturbation onset, relative to the values measured on the same individuals and perturbation conditions during Familiarization. Specifically, participants were asked to apply 100 *±* 40% the torque measured in Familiarization Yield during Task Yield, 200 *±* 40% the torque measured in Familiarization Yield during Task Resist, and 100 *±* 40% the torque measured in Familiarization Slow during Task Slow. Trials were considered successful if the measured torque fell within the prescribed range. If the trial was successful, the goal torque target turned green. If the trial was unsuccessful, the goal torque target remained red.

During the Resist condition in Familiarization sessions, visual feedback was provided after each perturbation, based on the average torque measured in the time comprised between 100 and 150 ms post-perturbation onset. This time frame was chosen to include a 50 ms delay from the [50 100] ms time interval typically considered for LLRs for upper extremity muscles. The delay was set to 50 ms to account for the electromechanical delay and to avoid participant success due to a “voluntary” response (typically considered as EMG signal measured *>*100 ms). Participants were instructed to apply 200 *±* 40% the torque that was measured in the same time range during the Yield condition, delivered just prior to the Resist condition during Familiarization (Fig. 2).

At the beginning of each session, participants were asked to complete five 1 s isometric contractions in extension at 0.2 *±* 0.035 Nm for EMG normalization purposes.

### 2.5 MRI Procedures

Imaging was performed using a Siemens 3T Prisma MRI system with a 64-channel head coil. Functional scans were acquired across two task sessions lasting approximately 15 min each. Scans were collected using a multiband multi-echo gradient-echo echo planar imaging sequence (TR=2.2 s, TEs=13.4/39.5/65.6 ms, FA=90 degrees, MB factor=2, GRAPPA=2, voxel resolution=1.731×1.731×4 mm^3^, 44 slices, FOV=180 mm, matrix size 104×104). Voxel resolution was kept small in-plane and longer out-of-plane, with voxel alignment to the brainstem to account for the orientation of brainstem nuclei which are very small but oblong axially.

A structural T1-weighted multi-echo magnetization-prepared rapid acquisition with gradient echo (MPRAGE) scan (TR=2.17 s, TEs=1.69/3.55/5.41 ms, TI=1.16s, FA=7 degrees, FOV=256×256 mm^2^, and voxel resolution = 1×1×1 mm^3^) was acquired at the end of each fMRI experiment for registration and normalization to a common space. The three echo images were combined via root mean square.

### 2.6 Data Processing and Statistical Analysis

#### 2.6.1 EMG Data Processing

A standard processing pipeline was used to process EMG data: band-pass filtering (4th order Butterworth 20-250 Hz) to remove motion artifacts and high frequency noise, signal rectification, and low-pass filtering (4th order Butterworth 60 Hz) to extract the envelope of the EMG signal.

FCR EMG signal was normalized by dividing the processed signal by the average signal measured in the last 200 ms during background contraction before perturbation onset across all perturbation trials. ECU EMG signal was normalized by dividing the processed signal by the average signal measured in five 1 s isometric contractions in extension conducted at the beginning of each session.

LLRa for each trial was calculated as the mean EMG signal 50-100 ms post perturbation onset, as has been done in previous studies (Weinman et al., 2021; Nikonowicz and Sergi, 2024). Additionally, background amplitude (BKGDa) and short latency response amplitude (SLRa) were calculated as the mean EMG signal 200-0 ms pre-perturbation onset and 25-50 ms post perturbation onset, respectively. SLRa was included as an outcome because it is understood to be a spinal reflex and thus should not be affected by supraspinal feedback incorporating task goals.

#### 2.6.2 Torque Data Processing

Torque data were processed using Matlab 2023a (Mathworks, Inc. Natick, MA, USA). For each perturbation trial, torque response amplitude was calculated as the mean torque measured 100-150 ms post perturbation onset. Success rates for each instruction condition were defined as whether the trial was in the red or green condition as displayed by the visual feedback (Fig. 2).

#### 2.6.3 Statistical Analysis

EMG and torque data were analyzed at a group level using a linear mixed model with condition (Yield, Resist, Slow) as the main effect and participant and the interaction of participant and condition as random effects. For participant-level analysis, no random effects were included, and a one-way ANOVA was used instead. Post-hoc Tukey tests were used to test for significant differences between conditions. For post-hoc tests, the Cohen’s effect size for paired samples *d_z_* was computed as the difference of the means divided by the standard deviation of the difference. All statistical analyses were conducted using JMP 17 Pro (SAS Institute Inc., Cary, NC, USA).

#### 2.6.4 fMRI Data Processing

Analysis of fMRI data was performed using a custom pipeline that integrates functions from FSL (version 6.0.7.7) (Jenkinson et al., 2012), AFNI (version 24.3.06) (Cox, 1996), and SPM12 (Wellcome Department of Cognitive Neurology, London, UK) running on Matlab 2023a. Structural T1-weighted images for each participant were processed using FSL using bias field correction and brain extraction. Functional data were preprocessed using FSL and AFNI. Head motion realignment was estimated for the first echo data referencing the single band reference image taken at the beginning of the scan and then applied to all echo timeseries. All images were brain-extracted and distortion corrected before Tedana (version 24.0.2) was used to calculated a T^∗^-weighted combination of the three echo datasets, producing an optimally-combined timeseries. Multi-echo independent component analysis (ME-ICA) was performed on the optimally-combined dataset using Tedana and resulting components were manually classified as accepted (signal of interest) or rejected (noise) (Reddy et al., 2024b). 4mm spatial smoothing was applied to functional data for whole brain analyses, while smoothing was left out of the preprocessing pipeline for brainstem-specific analyses.

SPM12 was used for first (participant-level) and second level (group-level) analyses. Functional datasets were treated as blocked design datasets rather than event design datasets to avoid dependence on accurate transient hemodynamic response modeling as brainstem hemodynamics are different than that of cortical hemodynamics. Separate first level analyses were conducted to determine whole-brain and brainstem-specific activation. The same general linear model was used for both analyses but the model regressors were convolved with a standard hemodynamic response function for whole-brain analysis and a brainstem-specific hemodynamic response function for brainstem-specific analysis (Zonnino et al., 2021; Kim et al., 2022). The brainstem-specific hemodynamic response function peaks earlier and has a smaller, earlier undershoot than the standard hemodynamic response function, reflecting the faster and more transient blood oxygen level dependent response observed in brainstem nuclei compared with cortical regions. The neural response was modeled as:

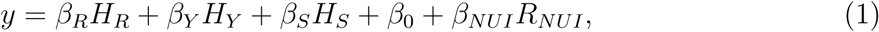

where *H_R_*, *H_Y_*, and *H_S_* are the task regressors (rectangular functions convolved with the hemodynamic response function) and *R_NUI_*is a set of 24+ nuisance regressors. Nuisance regressors included: 18 physiological noise regressors (6 cardiac, 8 respiratory, 4 interaction) and 6 head movement regressors calculated using the RETROICOR algorithm via the TAPAS PhysIO Toolbox (Kasper et al., 2017). Additionally, nuisance regressors included rejected regressors identified via multi-echo independent component analysis (ME-ICA) created using Tedana, as was done in Reddy et al. (2024b) and orthongonalized to all other regressors plus the accepted components from Tedana’s ICA, via AFNI’s 3dTproject (Medina et al., 2025). Note that unsuccessful trials were not accounted for by additional regressors.

General linear model estimation was used to identify voxels whose signal was significantly modulated by any experimental condition relative to rest (Slow*>*0: *β_S_ >* 0, Yield*>*0: *β_Y_ >* 0, Resist*>*0: *β_R_ >* 0), or significantly more in any experimental condition relative to the others (Resist*>*Yield: *β_R_− β_Y_ >* 0, Resist*>*Slow: *β_R_− β_S_ >* 0, Yield*>*Slow: *β_Y_− β_S_ >* 0). Then, participant-specific statistical parametric maps were used as inputs for second level group analysis. For brainstem-specific analysis, threshold-free cluster enhancement (TFCE) (Smith and Nichols, 2009) was performed within a mask of the brainstem created by thresholding the brainstem region of the Harvard-Oxford Subcortical Atlas (Makris et al., 2006; Frazier et al., 2005; Desikan et al., 2006; Goldstein et al., 2007) at 50%. Group-level analysis maps are reported using family-wise error correction (*α* < 0.05) and significance is set to *p <* 0.05. A more conventional, supplementary brainstem-specific analysis (figures available in Supplementary Materials) was conducted without ME-ICA rejected regressors and with a small-volume correction to control for the family-wise error rate (*α* < 0.05) in the same region of interest (Poldrack et al., 2008).

To break down the measured activation into clusters, SPM12’s *spm_clusters* was used on the resulting TFCE-corrected maps and clusters including more than 5 voxels were reported. Within these clusters, peaks of activation were identified automatically. Clusters were further broken down by anatomical region (Midbrain, Pons, and Medulla, defined based on the FreeSurfer Brainstem Substructures Atlas (Iglesias et al., 2015), thresholded at 50%), and by laterality (left or right). Within each anatomically-divided cluster, the three peaks with the greatest statistical parameters are reported for concise summary of cluster maxima, separately for each side.

To identify which nuclei may have contributed to the measured cluster activation, we compared the cluster maps to nuclei atlases from Brainstem Navigator (García-Gomar et al., 2022; Singh et al., 2021, 2020; García-Gomar et al., 2019; Bianciardi et al., 2018, 2015). For each cluster, we quantified spatial overlap with atlas-defined nuclei. A nucleus was considered a potential contributor if the overlapping voxels accounted for *≥*5% of the cluster volume. This threshold was used as a heuristic to identify nuclei with meaningful spatial overlap while avoiding attribution based on very small intersections between atlas labels and clusters. Qualitatively similar candidate nuclei were identified when modestly varying this threshold. Given the small size of many brainstem nuclei and the spatial resolution of our fMRI sequence, the atlas-based overlap analysis should be interpreted as identifying nuclei that are spatially consistent with the observed clusters rather than definitively localizing activation to specific structures.

We computed three outcomes to quantify laterality of the group-level activations in the brainstem at each axial slice (*z*=const). These metrics were designed to capture both the spatial distribution and magnitude across hemispheres and were conceptually derived from laterality index approaches commonly used in neuroimaging to quantify dominance of functional activation (Seghier, 2008). In all these calculations, we used data from all voxels in the ROI (defined as the union of Midbrain, Pons, Medulla from the FreeSurfer Atlas), with non-negative t-scores resulting from each of the three contrasts described above (R*>*0, R*>*Y, R*>*S). First, we computed the sum of *t* scores (*t_sum_*) for the left (*l*) and right (*r*) sides as an indicator of intensity and extent of activation from either side, as

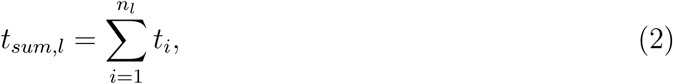

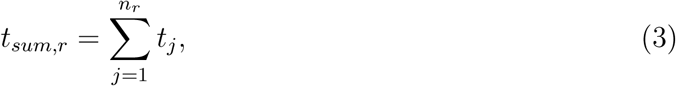

where *i* and *j* are indices corresponding to each voxel with non-negative score in a given slice, within the ROI, and *n_l_* and *n_r_* are the number of voxels with non-negative activation on the left and right sides, respectively. Our second outcome was the *t* score moment (*M*), which scales the contribution of each supra-threshold voxel proportionally to the lateral distance of each voxel from the midline (*x*), as

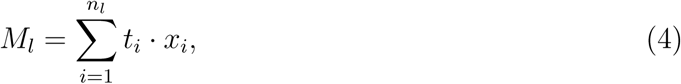

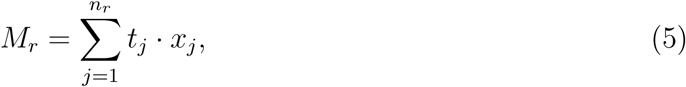

We then combined the outcomes above into the third outcome, which would indicate the lateral coordinate of the single voxel that would produce the same amount of moment imbalance for a given slice as the entire distribution of *t* scores in that slice. In analogy with the definition of center of mass in mechanical systems, we defined this outcome as center of activation (COA), with the same units as *x* (thus mm), which would be positive to indicate right dominance of the spatial distribution of t-scores, and negative otherwise.

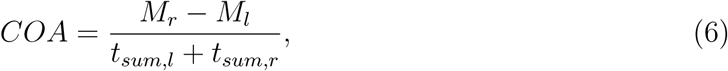

For whole-brain analysis, locations of significant activation were determined using FSL’s Juelich Histological Atlas, Harvard-Oxford Cortical Structural and Subcortical Atlases, and SUIT Probabilistic Cerebellum Atlas (Amunts and Zilles, 2020; Makris et al., 2006; Frazier et al., 2005; Desikan et al., 2006; Goldstein et al., 2007; Diedrichsen et al., 2009).

## 3 Results

### 3.1 Protocol Validation in the Mock Scanner

#### 3.1.1 Muscle Activity and Torque Performance during Perturbations

Group-level statistical analysis of FCR EMG data collected in Experiment 1 shows significant effects of instruction for all the outcomes considered (i.e. FCR activation during background, SLRa, and LLRa). LLRa in FCR (primary outcome) was greater for Resist compared to both Yield (Δ*LLRa_R_*_−_*_Y_* : 3.6141 *±* 1.9026, *p <* 0.001, *d_z_*=1.1773) and Slow (Δ*LLRa_R_*_−_*_S_*: 4.2140 *±* 1.7979, *p <* 0.001, *d_z_*=1.4527) conditions. Activity during Yield was not greater than Slow (Δ*LLRa_Y_* _−_*_S_*: 0.5999 *±* 0.3788, *p* = 0.7257, *d_z_*=0.9815), but on an individual participant level, LLRa was greater for Yield compared to Slow for 5*/*10 participants (Fig. 3). Background activity was greater for Yield compared to both Resist (Δ*BKGDa_Y_* _−_*_R_*: 0.1103 *±* 0.1126, *p* = 0.0063, *d_z_*=0.6069) and Slow (Δ*BKGDa_Y_* _−_*_S_*: 0.1522 *±* 0.0722, *p <* 0.001, *d_z_*=1.3066) conditions, but activity during Resist was not significantly different than during Slow (Δ*BKGDa_R_*_−_*_S_*: 0.0419 *±* 0.0987, *p* = 0.448, *d_z_*=0.2633). Importantly, the largest LLR responses occurred in the Resist condition despite similar background activation between Resist and Slow, suggesting that the observed LLR modulation reflects task-dependent feedback control rather than differences in baseline muscle activation.

**Figure 3:** (left) Group average of the FCR EMG measured in each perturbation condition across both task sessions in Experiment 1. The darker gray shaded region indicates time before perturbation onset and includes part of the background EMG activity. The lighter gray shaded region indicates the region of expected LLR activity. Thick lines represent the average and shaded areas represent *±*1 standard deviation (s.d.) indicating within-group variance. (right) Box plots showing the distribution of LLRa across participants with mean (horizontal white line), mean *±*1 standard error of the mean (dark shaded area), and mean *±*1 s.d. (light shaded area). Black dots indicate LLR mean of individual participants. Number of asterisks indicates significance level (\**p <* 0.05, \*\**p <* 0.005, \*\*\**p <* 0.001)

**Figure 4:**
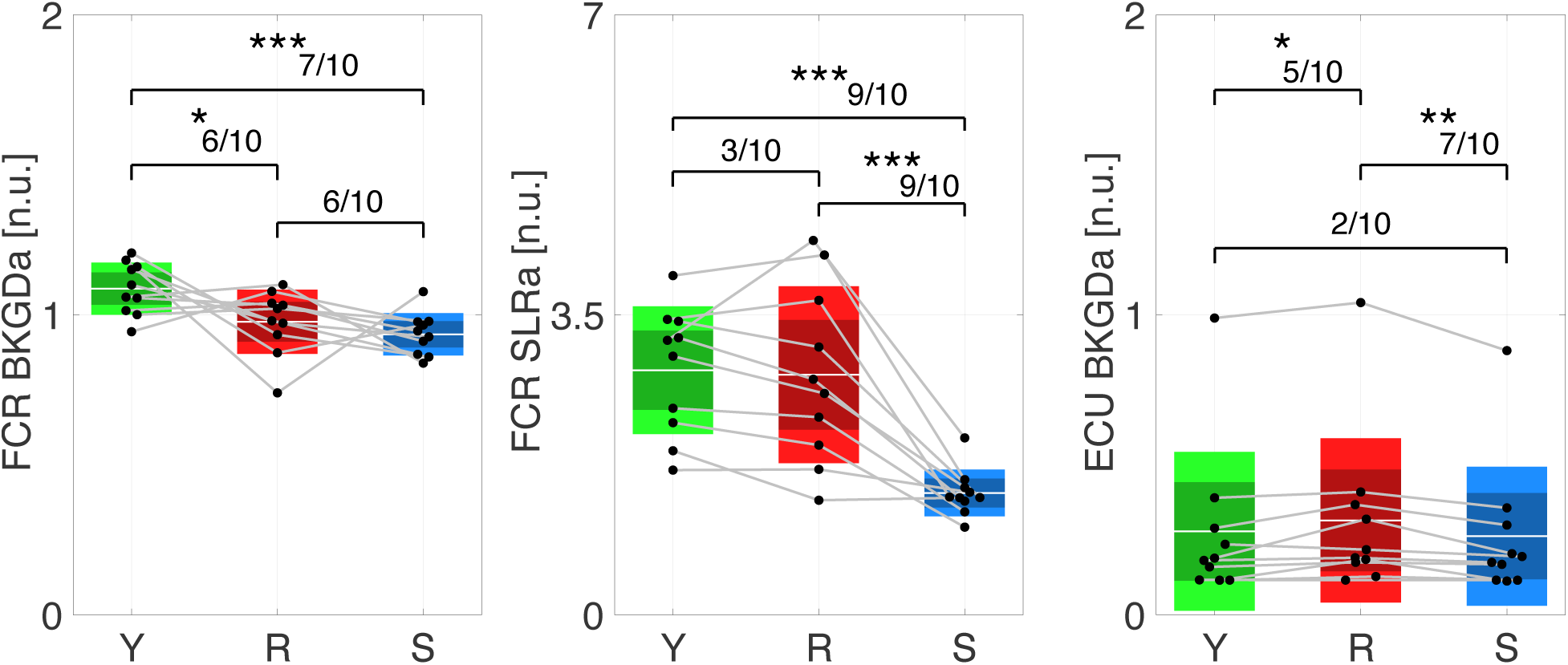
Secondary EMG Outcomes from Protocol Validation. As in Fig. 3 (right), all box plots show the distribution of average, normalized EMG across participants with mean (horizontal white line), mean *±*1 standard error of the mean (dark shaded area), mean *±*1 s.d. (light shaded area) and black dots indicate mean of individual participants. (left) Background FCR EMG. (center) Short latency response FCR EMG. (right) Background ECU EMG. Number of asterisks indicates significance level (\**p <* 0.05, \*\**p <* 0.005, \*\*\**p <* 0.001) the Yield and Slow conditions (Y: 99.63 *±* 0.92%, S: 100%), to lower levels of performance in the Resist condition (38.00 *±* 29.93%) (Fig. 6, left). Although Yield and Slow conditions did not consistently differ in reflex EMG responses, torque differed across all conditions. This difference likely reflects variation in wrist motion and passive mechanical interaction between the hand and the imposed perturbation rather than differences in voluntary muscle activation.

For the SLR time window, FCR activity was greater for Resist (Δ*SLRa_R_*_−_*_S_*: 1.3794 *±* 0.5440, *p <* 0.001, *d_z_*=1.5717) and Yield (Δ*SLRa_Y_* _−_*_S_*: 1.4320 *±* 0.4271, *p <* 0.001, *d_z_*=2.0780) compared to Slow, but not significantly different between Yield and Resist (Δ*SLRa_Y_* _−_*_R_*: 0.0526 *±* 0.3176, *p* = 0.9702, *d_z_*=0.1026). Detailed fixed effects results are available in Table 1.

**Table 1:** Mixed Model Results: Fixed Effects for All Validation Experiment Outcomes.

| **FCR RFXa** | **N par** | **DF Den** | **F Ratio** | **Prob>F** |
| --- | --- | --- | --- | --- |
| Instruction | 2 | 18.0 | 17.1209 | **<0.0001** |
| **FCR BKGDa** |  |  |  |  |
| Instruction | 2 | 54.7 | 10.4247 | **<0.0001** |
| **FCR SLRa** |  |  |  |  |
| Instruction | 2 | 18.0 | 26.2391 | **<0.0001** |
| **ECU BKGDa** |  |  |  |  |
| Instruction | 2 | 18.0 | 7.0611 | **0.0054** |
| **Avg. FB Torque** |  |  |  |  |
| Instruction | 2 | 18.0 | 136.2297 | **<0.0001** |

Group-level statistical analysis of background ECU EMG data shows a significant effect of instruction. Specifically, activity during Resist was greater compared to Slow (Δ*BKGDa_R_*_−_*_S_*: 0.0526 *±* 0.0326, *p* = 0.0048, *d_z_*=1.0008) and to Yield (Δ*BKGDa_R_*_−_*_Y_* : 0.0366 *±* 0.0279, *p* = 0.0499, *d_z_*=0.8135), but activity was not significantly different between Yield and Slow (Δ*BKGDa_Y_* _−_*_S_*: 0.0159 *±* 0.0231, *p* = 0.5199, *d_z_*=0.4279). This pattern is consistent with increased antagonist co-contraction during the Resist condition, which may reflect stabilization of the wrist in preparation to counteract the imposed perturbation.

Group-level statistical analysis of torque data shows a significant effect of instruction for torque measured in the feedback window (Fig. 5). Resist torque was larger than Yield (Δ*T_R_*_−_*_Y_* : 0.2646 *±* 0.0615, *p <* 0.001, *d_z_*=2.6662) and Slow (Δ*T_R_*_−_*_S_*: 0.4729 *±* 0.0673, *p <* 0.001, *d_z_*=4.3528) and Yield torque was larger than Slow (Δ*T_Y_* _−_*_S_*: 0.2083 *±* 0.0344, *p <* 0.001, *d_z_*=3.7493). These results were the same on an individual participant level for every participant. Significant effects on torque were measured across all conditions even though participants were not always able to modulate torque as cued by biofeedback. The average torque success rate varied between conditions, and ranged from almost perfect in

**Figure 5:**
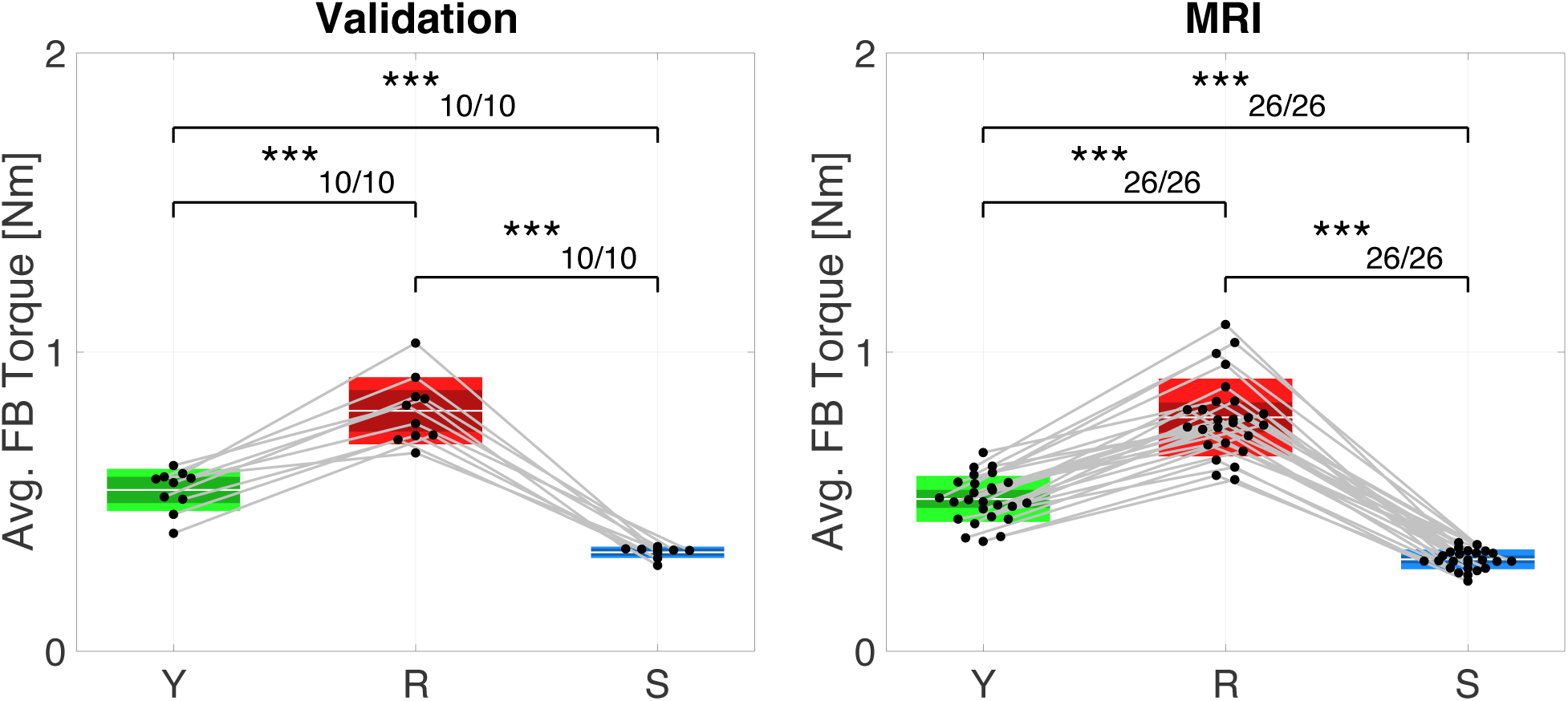
Average torque measured in the feedback window. (left) Validation experiment (right) MRI experiment. As in Fig. 3 (right), box plots show the distribution of average feedback window torque across participants with mean (horizontal white line), mean *±*1 standard error of the mean (dark shaded area), and mean *±*1 s.d. (light shaded area). Black dots indicate mean of individual participants. Number of asterisks indicates significance level (\**p <* 0.05, \*\**p <* 0.005, \*\*\**p <* 0.001)

Overall, these results indicate that task instruction selectively modulated LLRs, with greater LLRa during Resist compared to Yield and Slow, while SLR amplitudes were similar between Resist and Yield. These findings are consistent with prior work showing that goal-dependent modulation primarily affects the long-latency component of stretch responses.

### 3.2 Perturbations in the MRI Scanner

#### 3.2.1 Participant Torque Performance

In absence of in-scanner EMG data, torque was used as a proxy for task performance in the MRI experiment. Group-level statistical analysis of torque data shows a significant effect of instruction for torque measured in the feedback window, consistent with expectations. Consistent with the out-of-scanner validation experiment, Resist torque was larger than Yield (Δ*T_R_*_−_*_Y_* : 0.2723 *±* 0.0397, *p <* 0.001, *d_z_*=2.6392) and Slow (Δ*T_R_*_−_*_S_*: 0.4744 *±* 0.0438, *p <* 0.001, *d_z_*=4.1614) and Yield torque was larger than Slow (Δ*T_Y_* _−_*_S_*: 0.2021 *±* 0.0194, *p <* 0.001, *d_z_*=4.0133) (Fig. 5, right). These statistical differences between conditions were also true on an individual participant level for all participants. Success rates were also similar as in the out-of-scanner validation experiment, with an average torque response success rate of 95.10 *±* 5.71%, 50.29 *±* 29.76%, and 99.86 *±* 0.59% for Yield, Resist, and Slow conditions, respectively (Fig.6,5).

**Figure 6:**
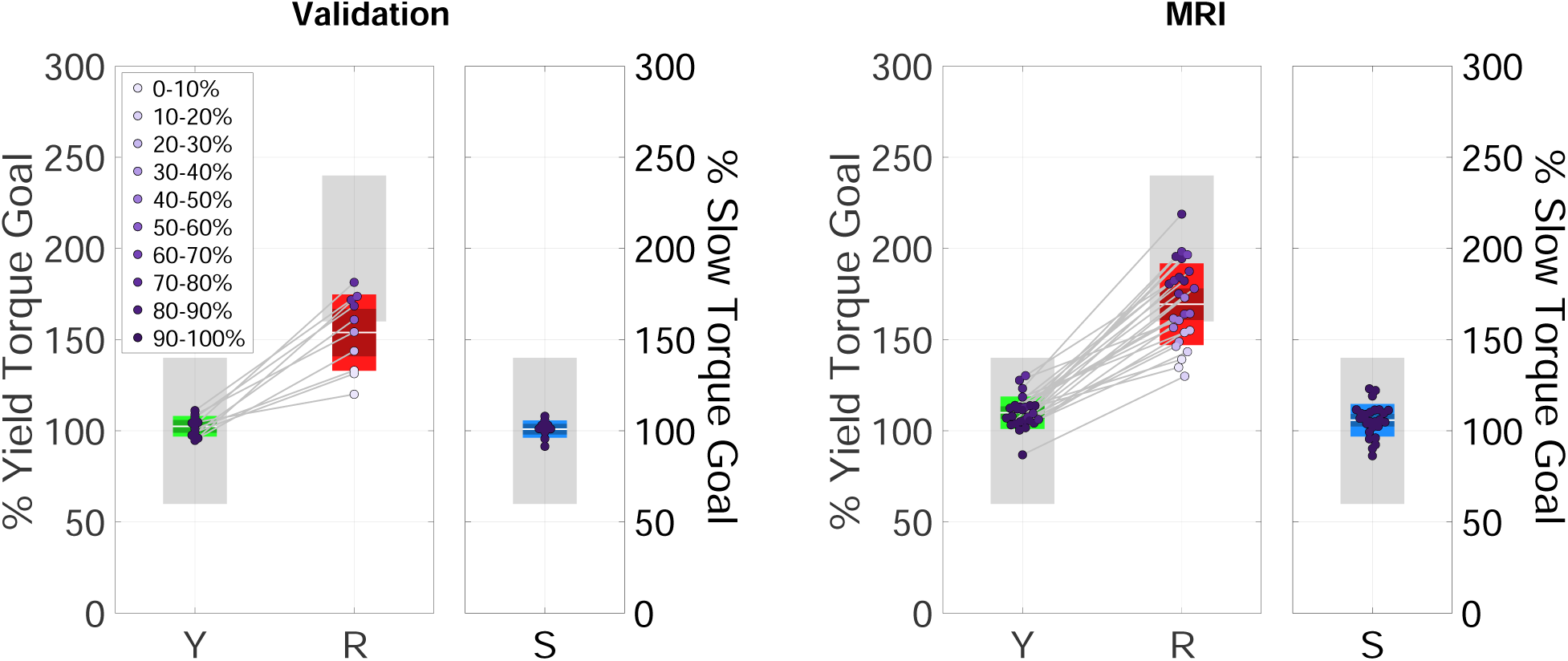
Average torque measured in the feedback window and normalized by torque goal with success rates for each individual participant. (left) Validation experiment (right) MRI experiment. As in Fig. 3 (right), box plots show the distribution of average feedback window torque across participants with mean (horizontal white line), mean *±*1 standard error of the mean (dark shaded area), and mean *±*1 s.d. (light shaded area). Dots indicate mean of individual participants with the color indicating success rate.

#### 3.2.2 Brainstem Activation

Brainstem-specific analysis revealed significant activity for Resist*>*0, Resist*>*Yield, and Resist*>*Slow contrasts. Tables 2, 3, and 4 include the top three peaks of activation on the left and right sides of the brainstem, broken down by cluster and brainstem region. Notably, the contrasts Slow*>*0, Yield*>*0, and Yield*>*Slow did not result in any significant activation in the brainstem.

**Table 2:**
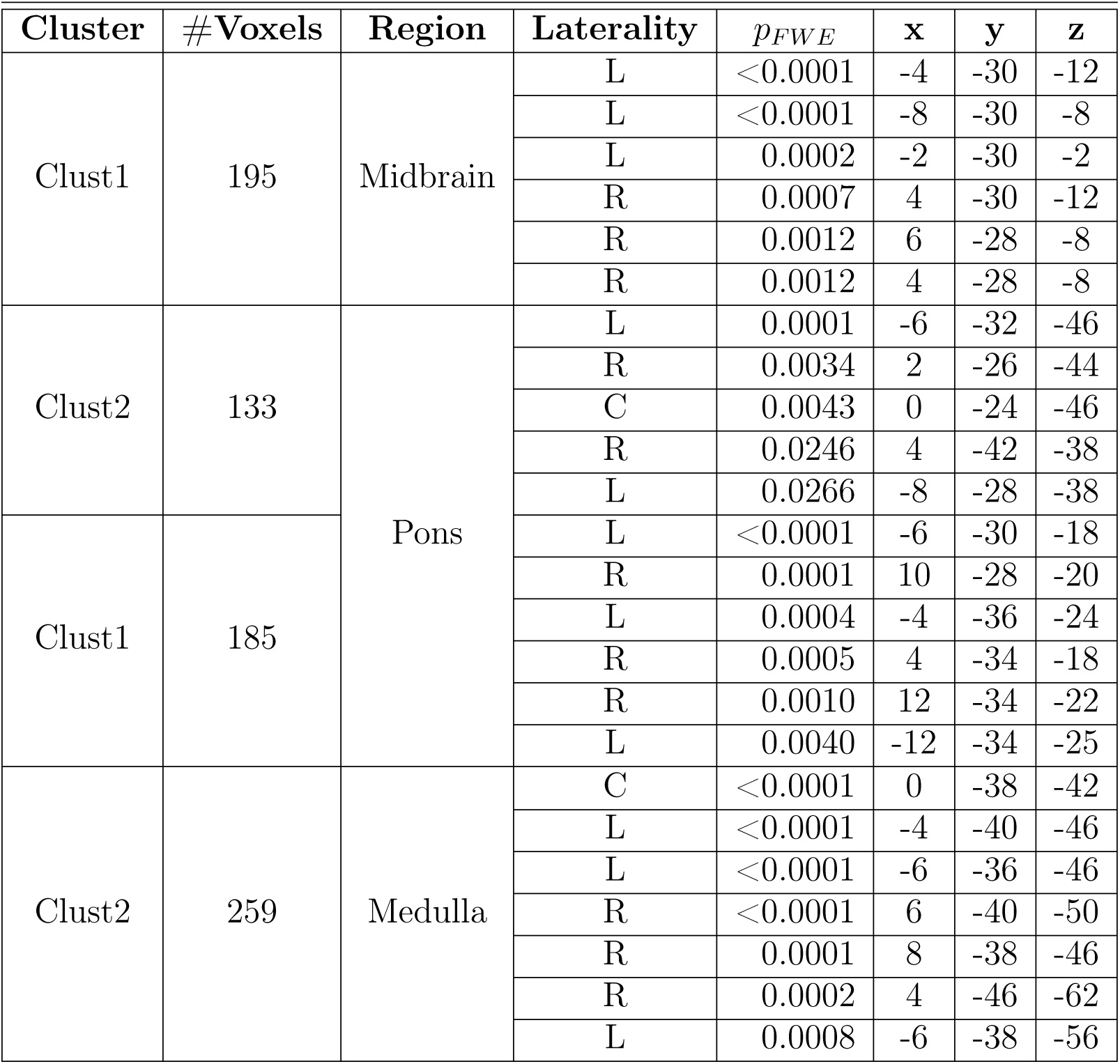
Brainstem Regions Activation: Resist*>*0.

**Table 3:** Brainstem Regions Activation: Resist*>*Yield.

| Cluster | #Voxels | Region | Laterality | $p_{FWE}$ | **x** | **y** | **z** |
| --- | --- | --- | --- | --- | --- | --- | --- |
| Clust1 | 95 | Midbrain | L | 0.0009 | -8 | -30 | -4 |
|  |  |  | L | 0.0010 | -2 | -30 | -2 |
|  |  |  | R | 0.0022 | 4 | -30 | -6 |
|  |  |  | L | 0.0022 | -6 | -34 | -4 |
|  |  |  | R | 0.0038 | 8 | -30 | -2 |
| Clust2 | 117 | Pons | L | 0.0005 | -8 | -42 | -40 |
|  |  |  | L | 0.0005 | -8 | -36 | -44 |
|  |  |  | L | 0.0008 | -4 | -40 | -38 |
|  |  |  | R | 0.0028 | 8 | -40 | -38 |
|  |  |  | R | 0.0029 | 8 | -34 | -46 |
|  |  |  | R | 0.0160 | 10 | -36 | -38 |
|  |  |  | C | 0.0233 | 0 | -28 | -44 |
| Clust1 | 174 |  | L | 0.0008 | -10 | -30 | -20 |
|  |  |  | L | 0.0010 | -6 | -36 | -24 |
|  |  |  | L | 0.0011 | -4 | -28 | -20 |
|  |  |  | R | 0.0030 | 12 | -34 | -22 |
|  |  |  | R | 0.0048 | 10 | -28 | -20 |
|  |  |  | R | 0.0048 | 4 | -34 | -26 |
| Clust2 | 174 | Medulla | C | <0.0001 | 0 | -38 | -42 |
|  |  |  | R | 0.0003 | 2 | -36 | -46 |
|  |  |  | R | 0.0004 | 8 | -38 | -46 |
|  |  |  | L | 0.0004 | -6 | -40 | -42 |
|  |  |  | R | 0.0007 | 4 | -44 | -60 |
|  |  |  | C | 0.0039 | 0 | -42 | -50 |
|  |  |  | L | 0.0052 | -4 | -40 | -56 |
|  |  |  | L | 0.0054 | -6 | -44 | -60 |

**Table 4:**
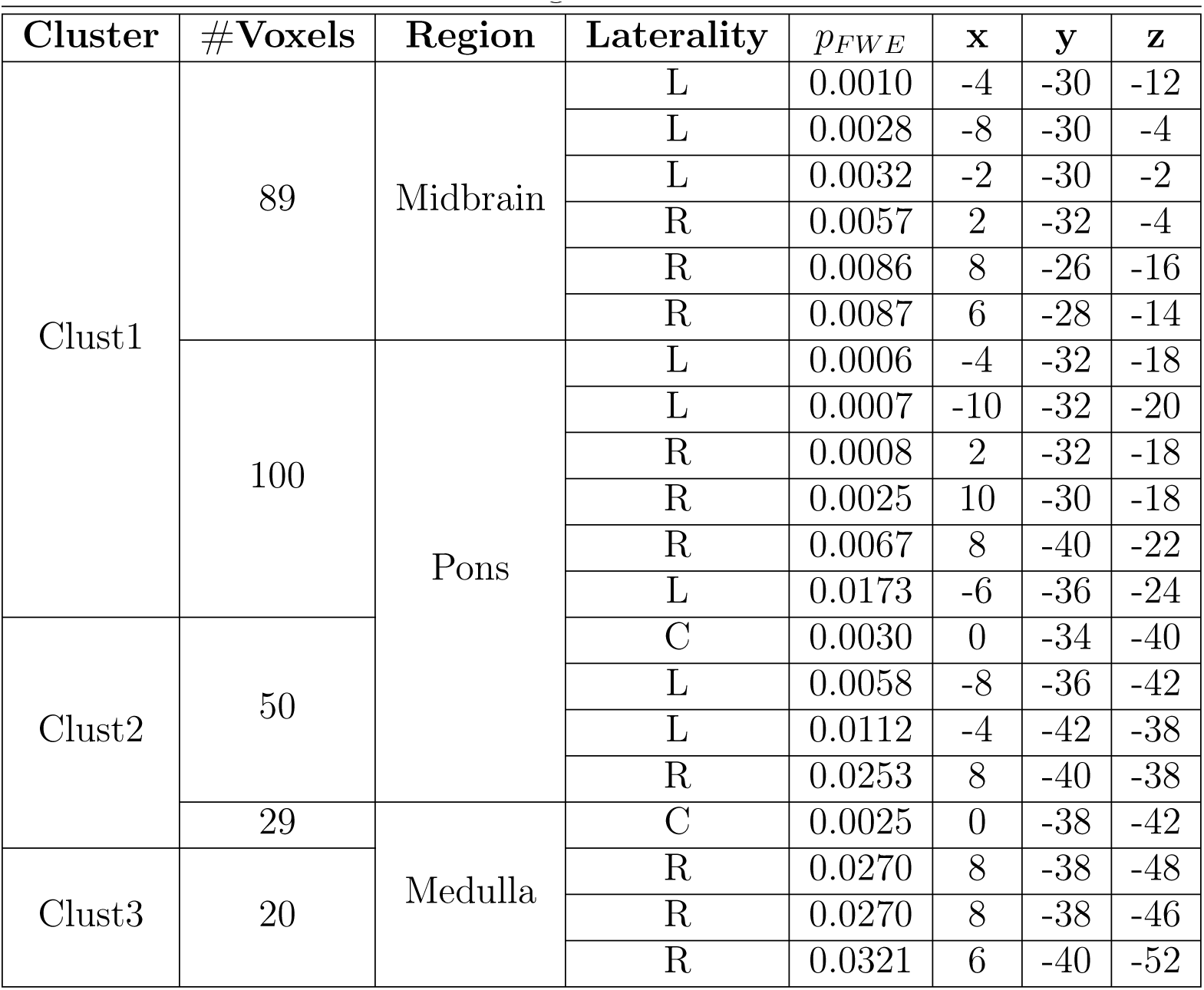
Brainstem Regions Activation: Resist*>*Slow.

As seen in Fig.7 and Table 2, significant activity for Resist*>*0 was widespread, with bilateral clusters spanning the midbrain, pons, and medulla. There were 2 clusters with more than 5 above-threshold voxels. The first cluster’s peaks spanned the midbrain and pons bilaterally, while the second cluster’s peaks spanned the pons and medulla bilaterally. The first cluster mostly overlapped bilaterally with the superior colliculi and periaqueductal gray and ipsilaterally with the pontine reticular nucleus. The second cluster mostly overlapped bilaterally with the lateral inferior medullary reticular formations, inferior olivary nuclei, and pontine reticular nuclei and contralaterally with the laterodorsal tegmental nucleus. A reference guide for nuclei locations is provided in Fig. 14.

**Figure 7:**
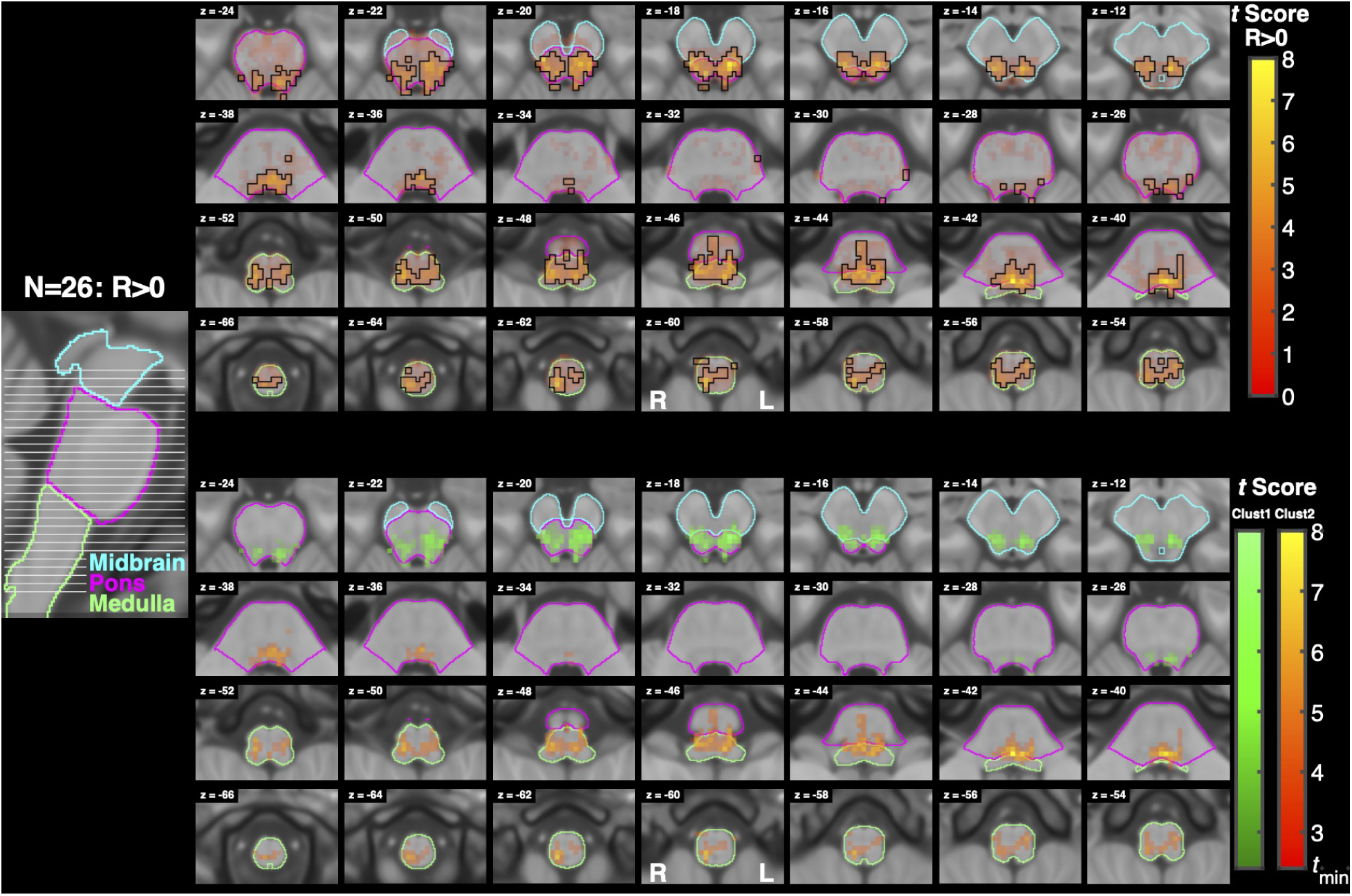
Activation associated with the Resist*>*0 contrast. A sagittal slice on the left shows the location of each axial slice of the brainstem shown on the right with the highest slice on the top right and the lowest slice on the bottom left. Three sections of the brainstem are outlined with different colors to facilitate orientation. (top) Functional maps for the contrast Resist*>*0. Voxel color maps to *t* score (no lower limit), while opacity maps to TFCE significance, with significant voxels outlined in black. (bottom) Cluster analysis of activation measured for the Resist*>*0 contrast. Different clusters of significant activation are color coded, and voxel color is modulated by cluster and *t* score. For this analysis *t_min_* = 2.4.

Resist*>*Yield (Fig.8) and Resist*>*Slow (Fig.9) activity spanned similar locations as Resist*>*0, and were also widespread, with clusters localizing bilaterally across the midbrain, pons, and medulla. The Resist*>*Yield contrast resulted in 2 clusters with more than 5 voxels. As with the Resist*>*0 contrast, the first Resist*>*Yield cluster’s peaks spanned the midbrain and pons bilaterally, while the second cluster’s peaks spanned the pons and medulla bilaterally. The first cluster overlapped bilaterally with the pontine reticular nuclei, superior colliculi, and the periaqueductal gray. The second cluster mostly overlapped bilaterally with the lateral inferior medullary reticular formations and pontine reticular nuclei, and contralaterally with the inferior olivary nucleus and parvicellular reticular nucleus.

**Figure 8:**
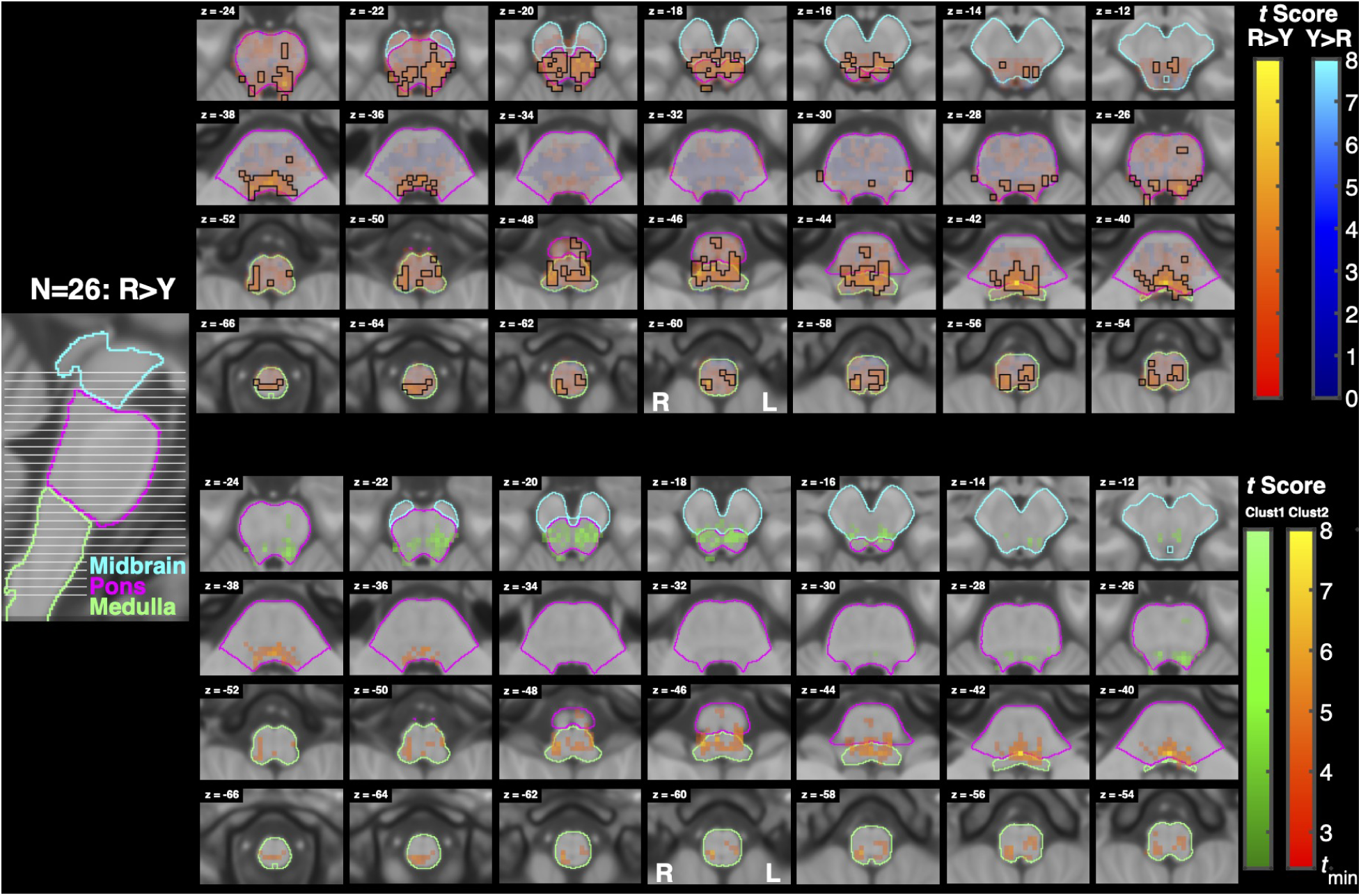
Activation associated with the Resist*>*Yield contrast. Conventions are the same as in Fig. 7. (top) Functional maps for the contrast Resist*>*Yield. (bottom) Cluster analysis of activation measured for the contrast Resist*>*Yield. For this analysis *t_min_* = 2.4.

**Figure 9:**
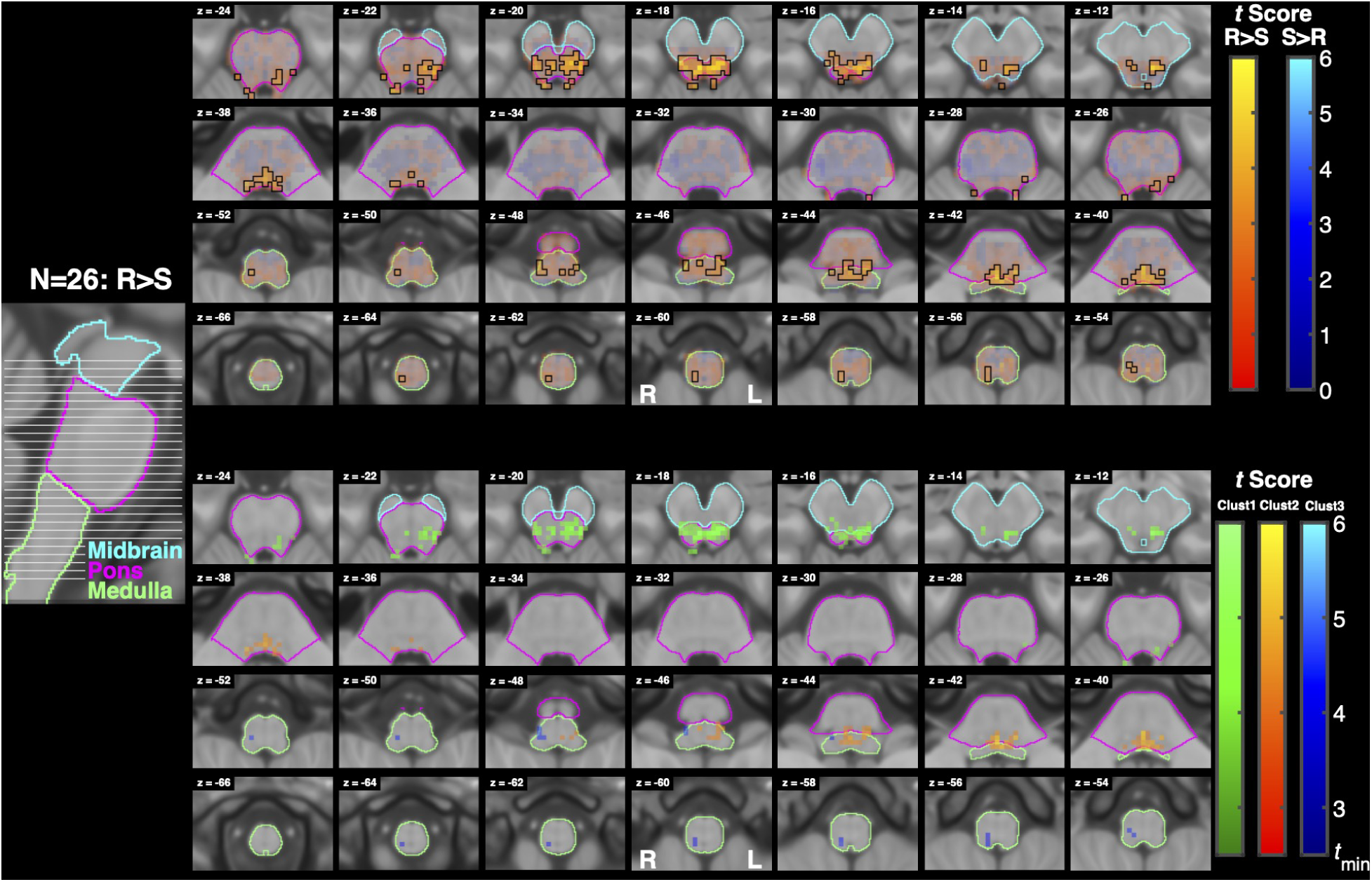
Activation associated with the Resist*>*Slow contrast. Conventions are the same as in Fig. 7. (top) Functional maps for the contrast Resist*>*Slow. (bottom) Cluster analysis of activation measured for the contrast Resist*>*Slow. For this analysis *t_min_* = 2.5.

The Resist*>*Slow contrast resulted in 3 clusters with more than 5 voxels. The first cluster’s peaks spanned the midbrain and pons bilaterally, the second cluster’s peaks spanned the pons and medulla bilaterally, and the third cluster’s peaks spanned the ipsilateral medulla. The first cluster overlapped with the periaqueductal gray, bilateral superior colliculi, and ipsilateral pontine reticular nucleus. The second cluster mostly overlapped bilaterally with the pontine reticular nuclei, laterodorsal tegmental nuclei, and vetibular nuclei complexes, contralaterally with the parvicellular reticular nucleus, subcoeruleus, superior olivary complex, and medial and lateral superior medullary reticular formations. Finally, the third cluster mostly overlapped ipsilaterally with the superior and inferior lateral medullary reticular formations, superior and inferior olivary nuclei, viscero-sensory-motor nuclei complex, inferior medial medullary reticular formation, and parvicellular reticular nucleus.

The results of our laterality analysis are summarized in Fig. 10 for all contrasts of interest (R*>*0, R*>*Y, R*>*S). All three outcomes of activation intensity (*t_sum_*), activation moment (*M*), and center of activation (COA) indicate a gradient of laterality of significant activation that changes from ipsilateral in medullar slices to contralateral in pontine slices, with no clear lateralization in the midbrain. Specifically, the center of activation is always lateralized ipsilaterally in medullar slices for all contrasts, while almost always lateralized contralaterally in pontine slices (always for R*>*0 and R*>*S, in most slices for R*>*Y).

**Figure 10:**
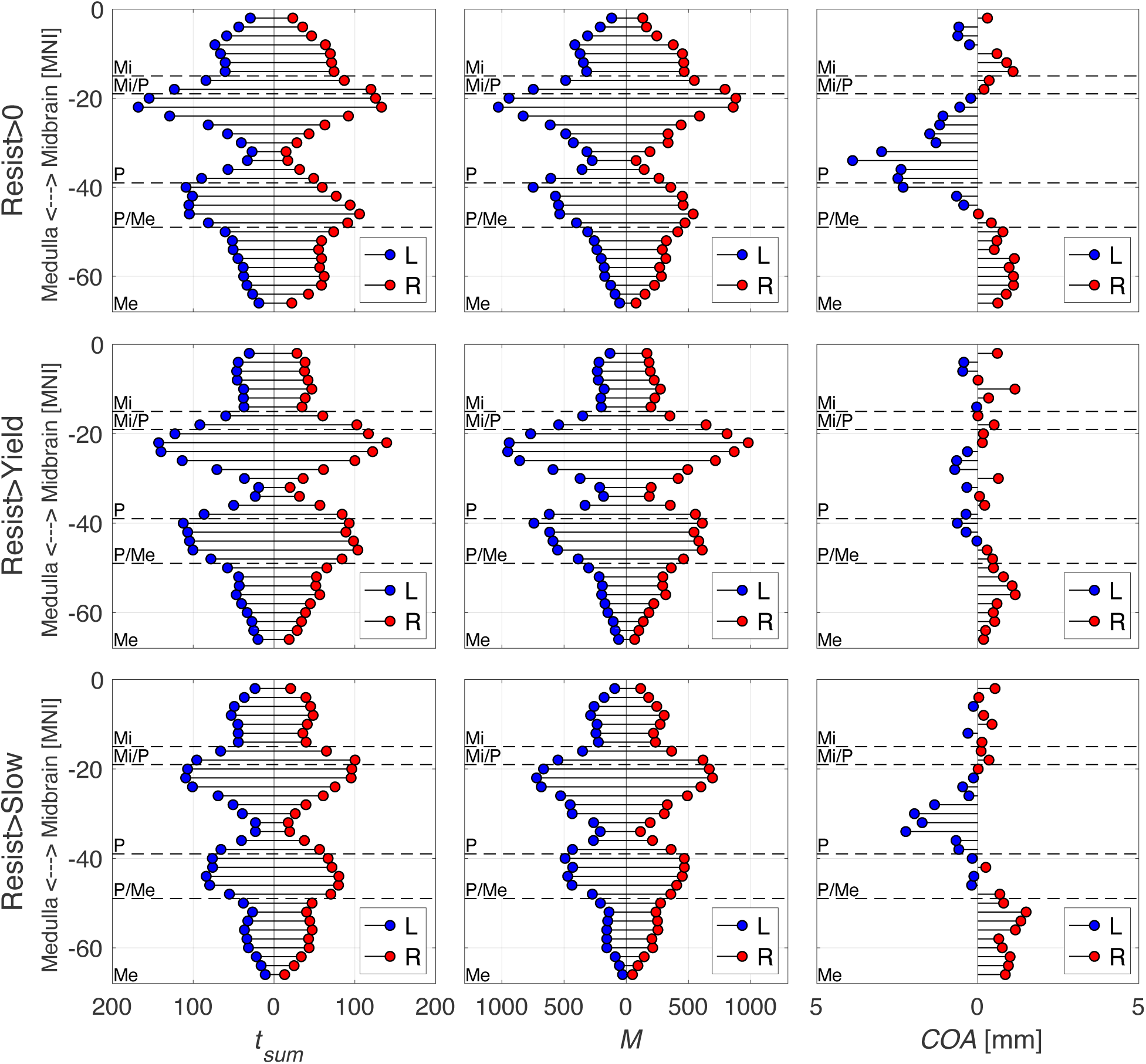
Laterality of positive activation in the brainstem. The three methods of quantifying laterality are shown with *t_sum_* on the left, *M* in the middle, and *COA* on the right with the contrasts Resist*>*0 on the top, Resist*>*Yield in the middle, and Resist*>*Slow on the bottom. Outcome values are on each x axis and axial slice location (*z* slice in MNI coordinates) are on each y axis. Horizontal dashed lines separate different sections of the brainstem by medulla (Me), pons (P), midbrain (Mi), and their combinations. Blue dots indicate activity from the left (contralateral) hemisphere and red dots indicate activity from the right (ipsilateral) hemisphere. Right is ipsilateral and left is contralateral.

**Figure 11:**
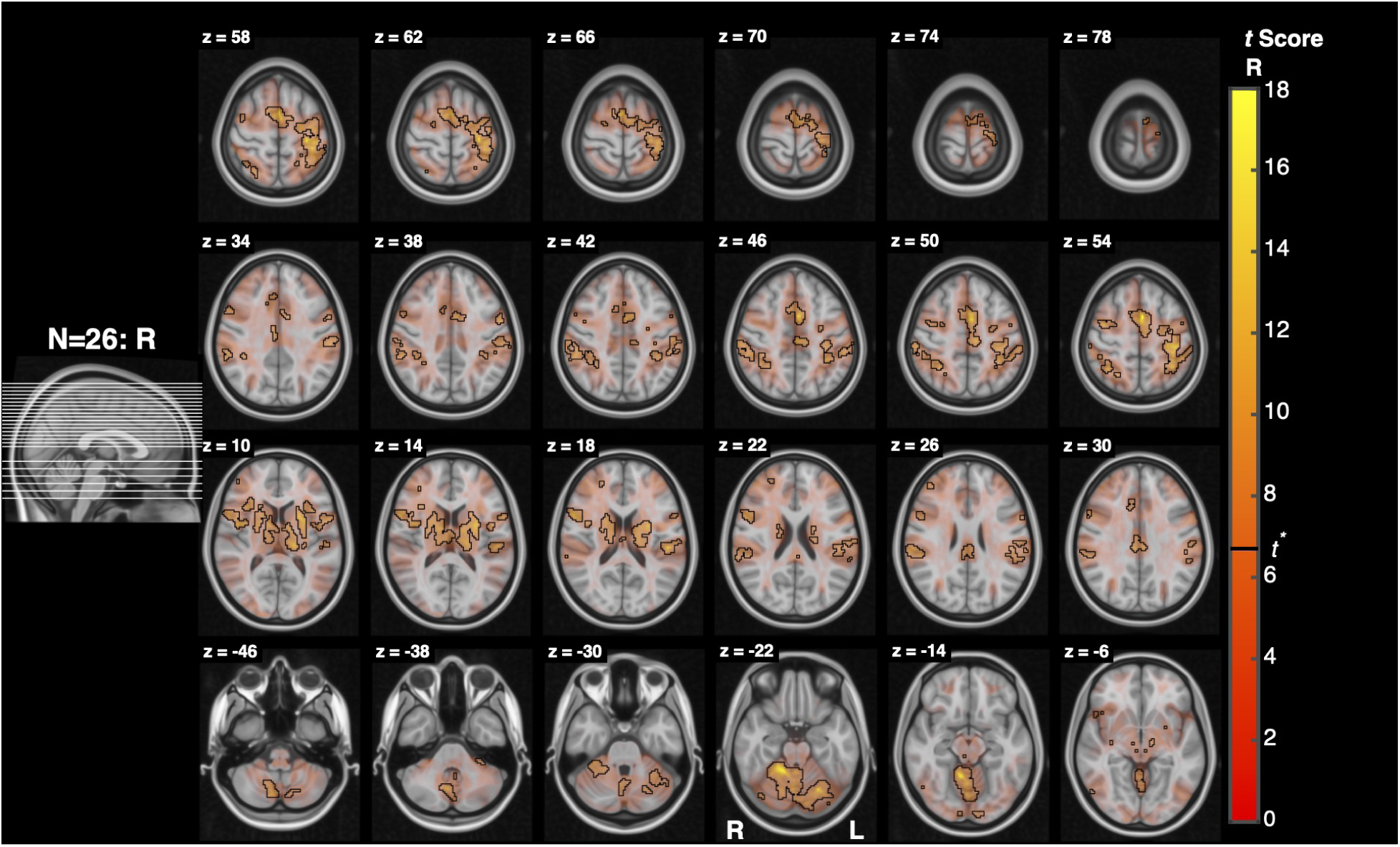
Cortical activation associated with the Resist*>*0 contrast. A sagittal slice on the left shows the location of each axial slice of the brain shown on the right with the highest slice on the top right and the lowest slice on the bottom left. Functional activation is shown with lower t score voxels in red and higher t score voxels in yellow. Opacity is also modulated by t score and significant voxels are outlined in black.

**Figure 12:**
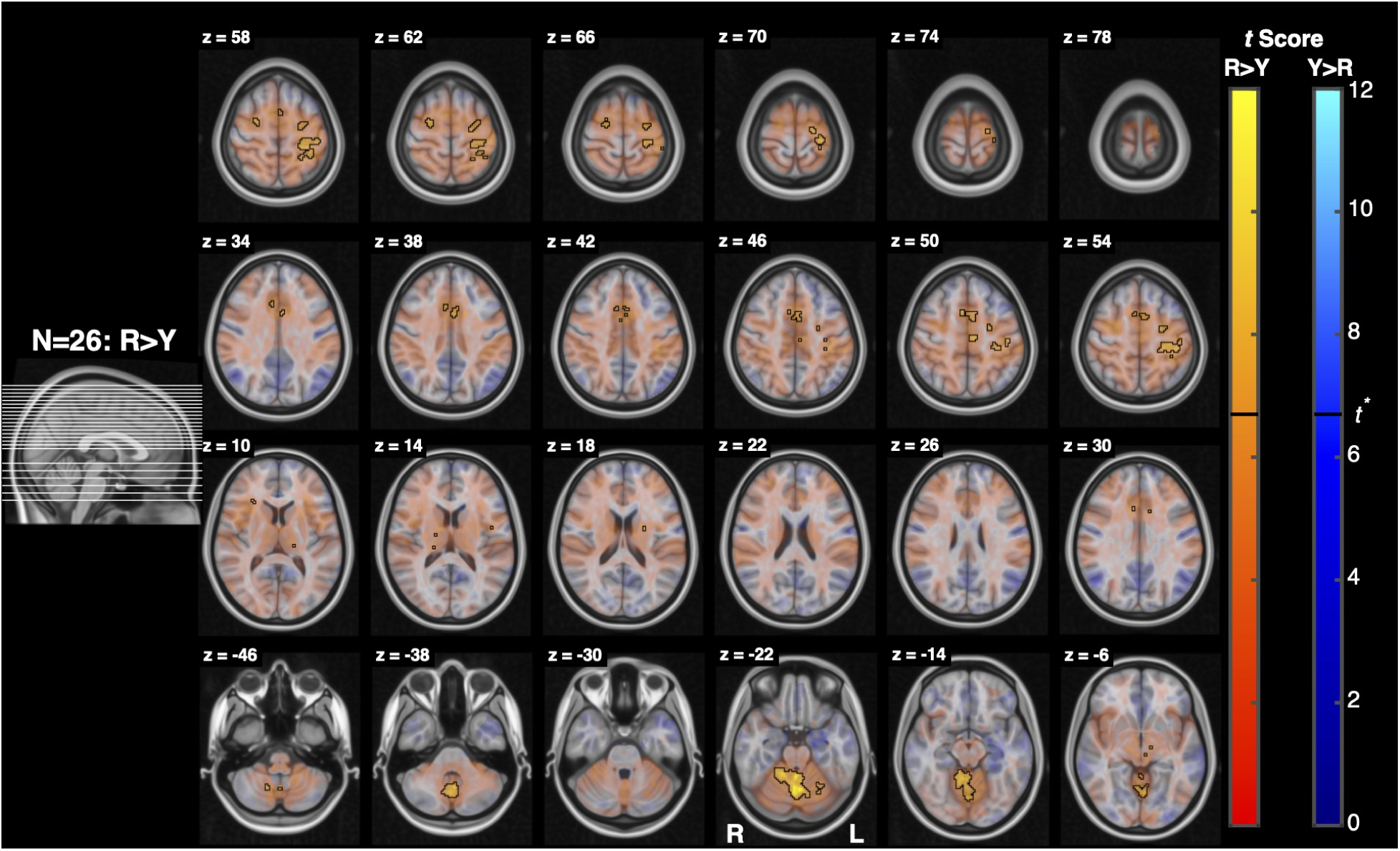
Cortical activation associated with the Resist*>*Yield contrast. Conventions are the same as in Fig. 11.

**Figure 13:**
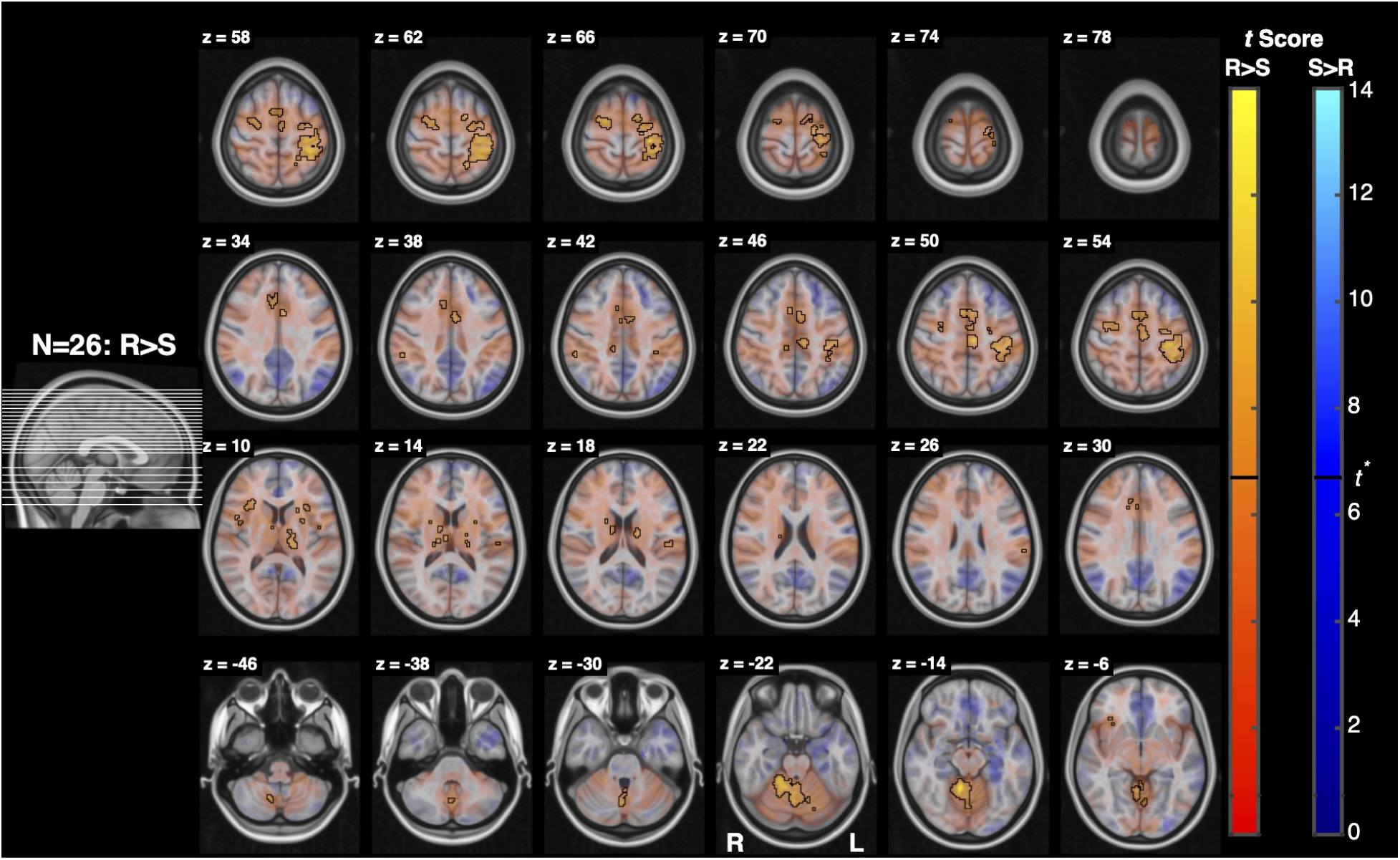
Cortical activation associated with the Resist*>*Slow contrast. Conventions are the same as in Fig. 11.

**Figure 14:**
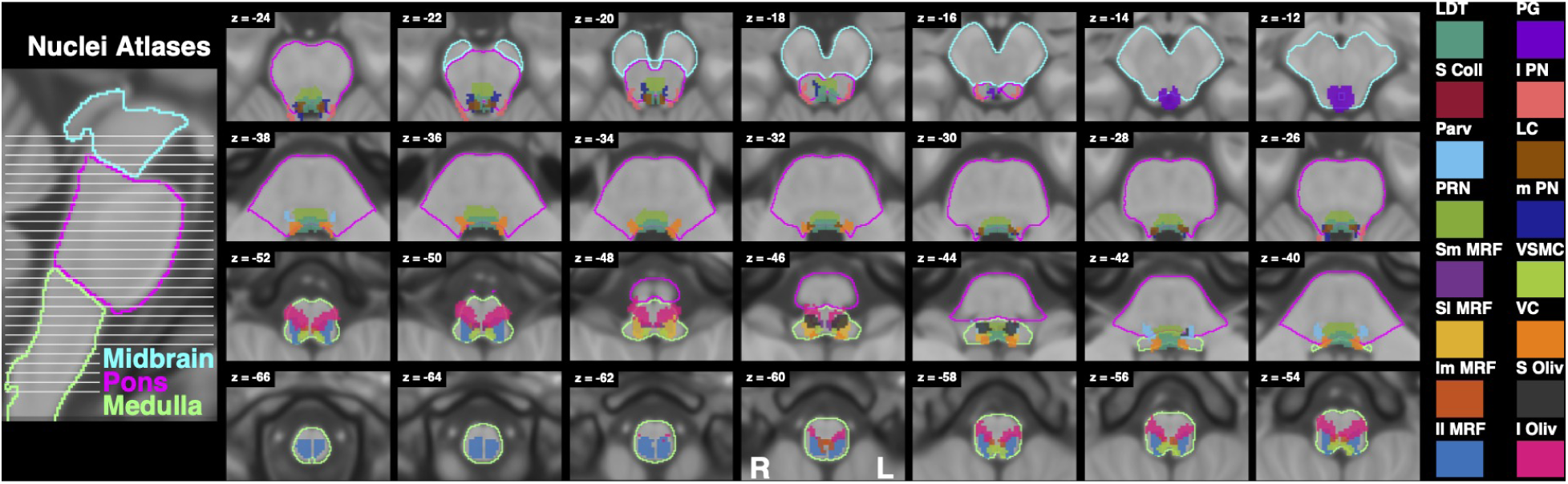
Brainstem slices with binarized atlases of referenced nuclei for comparison to functional activation figures. Abbreviations: laterodorsal tegmental nucleus (LDT), superior colliculus (S Coll), parvicellular reticular nucleus (Parv), pontine reticular nucleus (PRN), superior medial medullary reticular formation (Sm MRF), superior lateral medullary reticular formation (Sl MRF), inferior medial medullary reticular formation (Im MRF), inferior lateral medullary reticular formation (Il MRF), periaqueductal gray (PG), lateral parabrachial nucleus (l PN), locus coeruleus (LC), medial parabrachail nucleus (m PN), viscerosensorimotor nuclei complex (VSMC), vestibular nuclei complex (VC), superior olivary complex (S Oliv), and inferior olivary nucleus (I Oliv). Conventions are the same as in Fig. 7.

#### 3.2.3 Cortical Activation

Whole brain analysis revealed significant cortical activation for Slow*>*0, Yield*>*0, Resist*>*0, Resist*>*Yield, and Resist*>*Slow contrasts.

Activation associated with the contrast Slow*>*0 localized in the contralateral primary and secondary somatosensory cortices (S1, S2), premotor cortex (PM), and visual cortex (Brodmann Area 18). Additionally significant peaks localized in the ipsilateral visual cortex (Brodmann area 17), inferior temporal gyrus, posterior supramarginal gyrus, and inferior parietal lobule. Finally, bilateral peaks localized in the primary motor cortices, inferior lateral occipital cortices, and superior parietal lobules.

Yield*>*0 significant peaks were observed in the contralateral premotor cortex, primary motor cortex, primary and secondary somatosensory cortices, superior parietal lobule, visual cortex (Brodmann Area 18), insular cortex, and Broca’s Area 44, as well as the ipsilateral visual cortex (Brodmann Area 17) and anterior intraparietal sulcus hIP3.

Significant Resist*>*0 peaks in the contralateral visual cortex (Brodmann Area 18), ipsilateral Broca’s Area 44, central opercular cortex, anterior intraparietal sulci HIP1 and hIP3, visual cortex V1, anterior supramarginal gyrus, parietal operculum cortex, superior parietal lobule 7A, frontal pole, paracingulate gyrus, and occipital pole, as well as bilateral peaks in the premotor cortices, visual cortex V5, and posterior cingulate gyri.

Resist*>*Yield had significant peaks in the contralateral primary motor cortex, and primary somatosensory cortex, and bilateral anterior cingulate gyri, premotor cortices, and superior frontal gyri.

Resist*>*Slow had significant contralateral peaks in the primary and secondary somatosensory cortices, anterior cingulate gyrus, and superior frontal gyrus. Ipsilateral peaks localized in the premotor cortex, frontal orbital cortex, frontal operculum cortex, central opercular cortex, posterior supramarginal gyrus, and posterior cingulate gyrus. Bilateral activity localized in the primary motor cortices, insular cortices, and Broca’s area 44.

#### 3.2.4 Subcortical and Cerebellar Activation

Whole brain analysis revealed significant subcortical activation for Slow*>*0, Yield*>*0, Resist*>*0, Resist*>*Yield, and Resist*>*Slow contrasts.

Significant peaks for Slow*>*0 localized contralaterally in the thalamus and Crus I of the cerebellum, ipsilaterally in cerebellar lobes V, VIIb, and VIIIb, and bilaterally in the putamen and cerebellar lobe VI, as well as more centrally in lobe VI of the cerebellar vermis.

Yield*>*0 had significant contralateral peaks in the thalamus and Crus I of the cerebellum and ipsilateral peaks in cerebellar lobes I-IV, VI, VIIb, VIIIa, and VIIIb.

Resist*>*0 had significant contralateral peaks in the thalamus, putamen, and cerebellar lobes VI, VIIb, and X, ipsilateral peaks in cerebellar lobe V and Crus I, and bilateral peaks in cerebellar lobe VIIIa.

Resist*>*Yield had significant contralateral peaks in the thalamus and cerebellar lobes V and VI, ipsilateral peaks in cerebellar lobes I-IV and V, as well as more centrally located peaks in lobes VIIIa and IX of the cerebellar vermis.

Finally, Resist*>*Slow had significant contralateral peaks in the putamen and cerebellar lobe VI, ipsilateral peaks in cerebellar lobes V and VIIIb, and bilateral peaks in the thalamus.

## 4 Discussion

This study sought to determine whether human brainstem activity, particularly within reticulospinal nuclei, contributes to task-dependent modulation of stretch-evoked motor responses. Specifically, we tested whether instructing participants to resist a wrist perturbation would increase reticular formation activation relative to yielding, and whether the spatial organization of that activation would exhibit a rostrocaudal laterality gradient consistent with the double reciprocal model of reticulospinal control. Using high-resolution whole-brain fMRI optimized for the brainstem combined with a perturbation paradigm known to modulate long-latency responses, we quantified differences in BOLD activity under separate instructions and assessed laterality across brainstem slices.

### 4.1 H1: Task-Dependent Modulation of Brainstem Activity

Our first hypothesis predicts that stretch-evoked activity in the RF will be greater when participants are instructed to “resist” a perturbation than when instructed to “yield”. When resisting a perturbation, feedback gains are upregulated and prior work suggests that this increase cannot be fully explained by spinal or cortical circuits alone and instead, it reflects descending control from subcortical structures including the RST (Shemmell et al., 2009; Ravichandran et al., 2013). The current results support this hypothesis: the Resist*>*Yield contrast (Fig. 8) revealed significant bilateral activation throughout the pons and medulla, including regions consistent with the pontine and medullary reticular nuclei, inferior olivary nucleus, and parvicellular reticular nucleus. These areas largely overlap with nuclei that are considered primary motor origins of the RST, supporting the interpretation that task-dependent modulation of feedback responses involves reticulospinal recruitment (Peterson, 1979; Pruszynski and Scott, 2012; Kurtzer, 2014).

The specific nuclei implicated in the Resist*>*Yield contrast have well-established roles in postural control, sensorimotor integration, and muscle tone regulation. The pontine reticular, gigantocellular and parvicellular reticular nuclei have well-established origins of reticulospinal projections involved in regulating proximal limb muscle activity during reaching (Schepens and Drew, 2004; Riddle et al., 2009; Fisher et al., 2021). Additionally, although classically associated with cerebellar-driven motor learning, the inferior olivary nucleus also participates in timing and error-detection processes that influence response modulation. Prior work suggests inferior olivary nucleus involvement in adapting feedback responses (Zeeuw et al., 1998; Mundorf et al., 2025; Llinás, 2009). The periaqueductal gray, although not traditionally considered a group of motor nuclei, regulates arousal, multisensory integration, and orienting behaviors. Its activation here may reflect modulation of readiness or gain setting during the resist instruction rather than direct motor output (Koutsikou et al., 2015; Zhang et al., 2024). Notably, the expected gradient of activation across Slow, Yield, and Resist conditions was less evident than anticipated. Both the Yield*>*0 and Slow*>*0 contrasts produced minimal brainstem activation, resulting in Resist*>*Yield and Resist*>*Slow contrasts that were similar in spatial extent and magnitude (Figs. 8, 9). This observation suggests that task instructions may engage the brainstem more categorically than continuously. Rather than scaling linearly with effort or compliance, the transition to a “resist” task instruction may engage a relatively full reticulospinal command, with limited intermediate differentiation between Yield and Slow conditions. Under this interpretation, the Resist condition recruits a broader descending command to upregulate feedback gains, whereas Yield and Slow may rely more heavily on automatic sensorimotor feedback processes mediated cortically through long-latency reflex pathways, with comparatively limited additional brainstem involvement (Kurtzer, 2014; Perenboom et al., 2015).

Alternatively, the Yield condition may have relied more heavily on spinal feedback mechanisms with minimal supraspinal involvement leading to weaker brainstem differentiation between Slow and Yield conditions. Our EMG experiment resulted in LLR amplitudes that were not statistically different between Slow and Yield conditions, though there was a qualitatively notable change during the LLR window. Thus, it is possible that either our study was methodologically insensitive or our lack of a strong Yield*>*Slow contrast could reflect reduced RST contribution in those conditions.

The behavioral and EMG findings support this interpretation. Although torque production followed the expected ordering (Resist*>*Yield*>*Slow), the distinction between Yield and Slow was modest. In particular, LLRa was not significantly different between Yield and Slow, whereas Resist selectively enhanced LLRa relative to Yield. Moreover, SLRa showed clearer modulation between Yield and Slow, suggesting that spinal mechanisms may account for much of the differentiation between these two conditions. Together, these findings align with the imaging results, indicating that substantial reticulospinal engagement may be preferentially associated with the Resist instruction.

These findings differ in part with our previous StretchfMRI study (Zonnino et al., 2021), which reported significant activation in the medullary and pontine reticular formation during perturbations performed with a “yield” instruction. While the present study similarly detected stretch-related activity in the brainstem, we did not observe significant activation for the Yield condition alone (see Supplementary Material Fig. **??**).

Methodological differences from our earlier StretchfMRI study may help explain the discrepancy. Critically, Zonnino et al. modeled brainstem activity as scaling with trial-by-trial EMG amplitude, identifying regions in which BOLD signal covaried with the magnitude of the evoked muscle response. In contrast, the present study examined condition-level main effects relative to baseline, without explicitly testing for parametric relationships with EMG. These analytic approaches probe related but distinct questions: one tests whether brainstem activity tracks the magnitude of motor output, whereas the other tests whether overall activation differs between task instructions. Additionally, the prior study employed an event-related design rather than a block design, which may have differentially influenced habituation or the separation of stretch-related and non-stretch-related activity. The present study also incorporated physiological noise regression and multi-echo denoising procedures, which may have reduced false positives in small subcortical regions and yielded a more conservative estimate of stretch-related activation.

### 4.2 H2: Brainstem activations show a rostrocaudal laterlization gradient that is consistent with the double reciprocal model of brainstem motor control

Our findings provide compelling evidence that task instruction modulates activity in reticulospinal nuclei, underscoring the brainstem’s role as a dynamic contributor to goal-dependent feedback control. The bilateral distribution of activation across the pons and medulla sug- gests that both medial and lateral RST pathways are involved, consistent with their known bilateral projections (Baker, 2011; Buford, 2009). Importantly, the spatial pattern of activation reveals a structured organization that aligns with aspects of the double reciprocal model of the RF while also highlighting complexities beyond a simple two-tract framework.

The double reciprocal model proposes complementary roles for the descending RST pathways: the medial RST provides excitatory drive to contralateral extensor muscles and inhibitory signal to contralateral flexor muscles, while the lateral RST provides excitatory drive to ipsilateral flexor muscles and inhibitory signal to ipsilateral extensor muscles (Peterson et al., 1978; Sakai et al., 2009; Davidson et al., 2004; Davidson and Buford, 2006; Davidson et al., 2007). Recordings in non-human primates suggest general organizational tendencies along rostrocaudal and mediolateral axes, although these studies also emphasize the heterogeneity and bilateral branching of reticulospinal neurons. In this context, the model provides a useful framework for interpreting large-scale trends in activation patterns. Our results align with aspects of this organization but also highlight the predominantly diffuse and bilateral nature of brainstem activity during goal-dependent feedback control.

Across all three contrasts with significant brainstem activity, laterality analyses revealed a consistent ipsilateral-leaning pattern within the medulla, shifting toward a more bilateral or slightly contralateral-leaning pattern at the pontomedullary interface, and ultimately transitioning to contralateral-dominant activity in the pons, particularly in the Resist*>*0 and Resist*>*Slow contrasts. This inferior-superior gradient mirrors the expected anatomical arrangement. The lateral RST, originating in the medulla, has stronger ipsilateral projections while the medial RST, originating in the pons, has more bilateral/contralateral influence. The fact that the medulla consistently shows ipsilateral-biased activation while the pons shifts contralateral supports the interpretation that these clusters correspond to distinct components of the reticulospinal system.

Importantly, the overall gradient is stable across methods for computing laterality. How- ever, only the Resist*>*Slow contrast demonstrated strong unilateral dominance in the medulla. The gradient is thus more of a modest directional trend superimposed on a fundamentally bilateral system. The magnitudes of these laterality effects are modest. The *COA* in the medulla is consistently ipsilateral across contrasts, but the displacement is small, typically on the order of 1–2 mm. Similarly, laterality assessed using *t_sum_* reveals only moderate asymmetries. Even in the most lateralized contrast (Resist*>*Slow), the ipsilateral *t_sum_*in medullary slices is at most 150% of the contralateral value, indicating a bias rather than unilateral dominance. Pontine slices show the opposite directional trend, but again with relatively small spatial shifts and magnitude differences. Thus, although the direction of the gradient is reproducible across laterality metrics (*t_sum_*, *M*, and *COA*), the absolute degree of hemispheric separation remains limited.

Thus, rather than providing strong support for the double reciprocal model, our findings suggest (1) spatial differentiation along the pons/medulla boundary is present, mirroring the anatomical origins of the medial and lateral RSTs, (2) laterality changes gradually, not categorically, indicating that both pathways are likely co-activated during goal-dependent feedback responses, and (3) functional specialization may exist, but in humans it likely operates within an integrated and bilateral framework.

Functionally, resisting a perturbation requires both flexor responses to counteract the imposed stretch and extensor responses to stabilize the limb, which likely explains the simultaneous engagement of medullary and pontine nuclei. The bilateral nature of the responses is also expected. The RST is known to support whole-body postural adjustments and primate reticulospinal neurons frequently show bilateral or branching projections (Davidson and Buford, 2006).

### 4.3 Whole Brain Imaging Results

Whole-brain analyses revealed robust activation across a distributed sensorimotor network for all task conditions, including M1, S1, S2, PM, and superior parietal regions. These areas are known to support proprioceptive processing and corrective motor responses during limb perturbations (Scott, 2004; Omrani et al., 2016), confirming that the paradigm reliably engaged core sensorimotor circuits.

Yield*>*0 and Resist*>*0 showed broader recruitment of premotor and parietal regions than Slow*>*0, consistent with the greater demands on sensorimotor integration when participants intentionally modulated their feedback responses (Pruszynski and Scott, 2012).

The contrasts Resist*>*Yield and Resist*>*Slow highlighted increased activation in PM, cingulate, and frontal opercular areas which are regions associated with effort, action monitoring, and rapid updating of motor commands (Rushworth et al., 2004). Stronger S1 and M1 responses in these contrasts align with the larger corrective torques and enhanced long-latency responses observed behaviorally and via EMG.

Subcortically, activation in cerebellar sensorimotor regions (lobules V, VI, VIII) increased significantly for all contrasts except Yield*>*Slow. These regions contribute to online error correction and the integration of proprioceptive information to update motor commands and shape long-latency feedback responses (Zonnino et al., 2021; Manto et al., 2012).

In addition to cerebellar activation, whole-brain analyses revealed putamen and thalamic involvement across task conditions. Putamen activation was observed bilaterally in the Slow*>*0 contrast and contralaterally in both Resist*>*0 and Resist*>*Slow contrasts, suggesting a contralateral-dominant but bilateral pattern consistent with its role in upper limb motor control. The putamen is a central component of cortico-basal ganglia circuits involved in the execution and parametric control of voluntary movement, including the regulation of movement scaling and vigor, and is known to integrate proprioceptive input to modulate on-going motor output rather than initiate it (Turner and Desmurget, 2010; DeLong and Strick, 1974; Shirinbayan et al., 2019). Its engagement across conditions is therefore consistent with sustained voluntary contraction demands and the modulation of motor output during interaction with external loads rather than condition-specific differences between Yield and Resist.

The thalamus, which exhibited predominantly contralateral activation across all task conditions and bilateral activation in the Resist*>*Slow contrast, serves as a key relay between both cerebello-thalamo-cortical and basal ganglia-thalamo-cortical circuits. During perturbation tasks, the thalamus plays a critical role in integrating and transforming proprioceptive and cerebellar error-related signals to support online motor corrections, with activation shown to track moment-by-moment position errors and reflect context-dependent gating of sensory input (Suminski et al., 2007; Butler et al., 1998; Bosch-Bouju et al., 2013). The observed pattern of thalamic activation is therefore consistent with increased feedback control demands during perturbation conditions while also supporting the relay of basal ganglia output during voluntary motor engagement.

Notably, Resist*>*Yield did not result in robust putamen or thalamic activation, suggesting that while these subcortical structures contribute to general motor execution and feedback integration across tasks, they are less directly involved in the task-dependent modulation of LLRs. This distinction supports the interpretation that such modulation may instead rely more heavily on brainstem pathways.

Finally, cortical maps for the Resist*>*Yield and Resist*>*Slow contrasts qualitatively, though not significantly, showed patterns consistent with reduced activation in regions typically associated with the default mode network, suggesting that the Resist condition shifts processing toward task-related demands and away from internally focused activation. This interpretation is consistent with our understanding that there is task-related suppression of default mode network activation during cognitively demanding tasks (Buckner et al., 2008).

### 4.4 Muscle Activity and Torque Performance

Experiment 1 confirmed that participants produced distinct neuromuscular responses across the three tasks. As expected, the tasks produced clearly different levels of torque: Resist generated the largest amount of torque, Yield produced less, and the Slow condition elicited a minimal response. This same ordering appeared in Experiment 2. These parallel behavioral outcomes suggest that participants successfully understood and executed the task under both experimental environments.

The EMG results collected in Experiment 1 further validate that the perturbation paradigm elicits instruction dependent modulation of LLRa, consistent with previous studies (Lewis et al., 2006; Pruszynski et al., 2008; Weiler et al., 2015). Importantly, Resist background activity was not larger than that of Yield or Slow, indicating that the greater torque produced during Resist relied more heavily on the fast feedback responses as opposed to elevated anticipatory muscle activation. The observed differences in SLRa and LLRa support this interpretation: both response epochs were facilitated in Resist compared to Slow and LLRa was selectively enhanced in Resist compared to Yield.

Importantly, ECU EMG showed a modest but statistically significant effect of instruction on background activity. Participants exhibited elevated ECU activity during Resist compared to both Yield and Slow conditions, which may reflect increased co-contraction when preparing for a more challenging task. Greater antagonist activation could contribute to increased joint impedance via changes in muscle short-range stiffness, representing a potential confound when interpreting task-dependent differences in evoked responses. However, the magnitude of this background modulation was smaller than the observed modulation of LLRa, suggesting that changes in anticipatory activation along are unlikely to account for the enhanced LLRa during Resist.

### 4.5 Limitations and Future Directions

Interpreting activity in the brainstem warrants caution. Brainstem fMRI is constrained by its small anatomical structures, high vascular density, and sensitivity to motion and physiological noise, all of which reduce spatial specificity even under optimized acquisition pipelines.

Although our design successfully isolated task-dependent differences between our three conditions, an important limitation is the absence of a no-perturbation or slow-perturbation control matched to Resist. As a result, we cannot fully dissociate neural activity related to the intention to resist from activity driven by the stretch-evoked feedback response itself. Both components likely contribute to the observed brainstem activation and future experiments with independent manipulation of intention and sensory input will be necessary to disentangle them.

Finally, we did not record forces from the non-perturbed (left) wrist. Without bilateral measurements, we cannot determine whether participants generated coordinated or symmetric torque responses, which could explain the bilateral functional imaging results. Future studies incorporating bilateral force sensors and a Resist condition with a slow perturbation would help clarify the specific contributions to intention, feedback, and bilateral coordination to task-dependent modulation.

## 5 Conclusion

This study demonstrates that goal-dependent modulation of rapid feedback responses is associated with measurable changes in human brainstem activity. Using an MRI-compatible robotic wrist perturbation paradigm combined with a brainstem-optimized multi-echo whole brain fMRI protocol, we identified bilateral activation within pontine and medullary regions consistent with major reticulospinal nuclei during a Resist instruction relative to Yield. Although activation patterns were largely diffuse and bilateral, laterality analyses revealed a modest rostrocaudal gradient consistent with known anatomical trends in reticulospinal organization. Importantly, behavioral and EMG results confirmed instruction-dependent modulation of long-latency responses, supporting the interpretation that descending pathways contribute to the observed imaging differences.

The absence of a clear activation gradient across Slow, Yield, and Resist conditions suggests that reticulospinal engagement may operate in a relatively categorical manner once a resistive motor set is adopted, rather than scaling linearly with task demands. At the same time, the diffuse and bilateral nature of the activation patterns underscores the distributed and multisynaptic organization of the human brainstem motor system.

More broadly, these findings establish the feasibility of integrating MRI-compatible robotics, physiological monitoring, and advanced denoising strategies to probe brainstem motor circuits noninvasively. This approach provides a foundation for future studies investigating descending control in health and disease, including stroke, spinal cord injury, and motor rehabilitation, where reticulospinal pathways are thought to play a critical compensatory role.

## 6 Ethics Statement

This data was collected in accordance with the Declaration of Helsinki and approved by the Investigation Review Board of the University of Delaware Protocol no. 1097082-11. sectionAuthor Contributions R.C.N.: conceptualization, data curation, formal analysis, investigation, methodology, resources, software, validation, visualization, writing (original draft, review and editing), visualization. N.A.R.: methodology, resources, software, writing (review and editing). M.C.M. methodology, resources, software, writing (review and editing). M.G.B.: resources, supervision, writing (review and editing). F.S. conceptualization, funding acquisition, investigation, methodology, project administration, supervision, writing (review and editing).

## Acknowledgments

The authors would like to acknowledge Cody A. Helm and Kristin Schmidt for their assistance with data collections.

