## Supplemental Figures for "Stretch-evoked motor responses in the brainstem are modulated by task instructions": Nikonowicz_YRpaper_bioRxiv_supp.pdf

### 1 Supplementary Materials

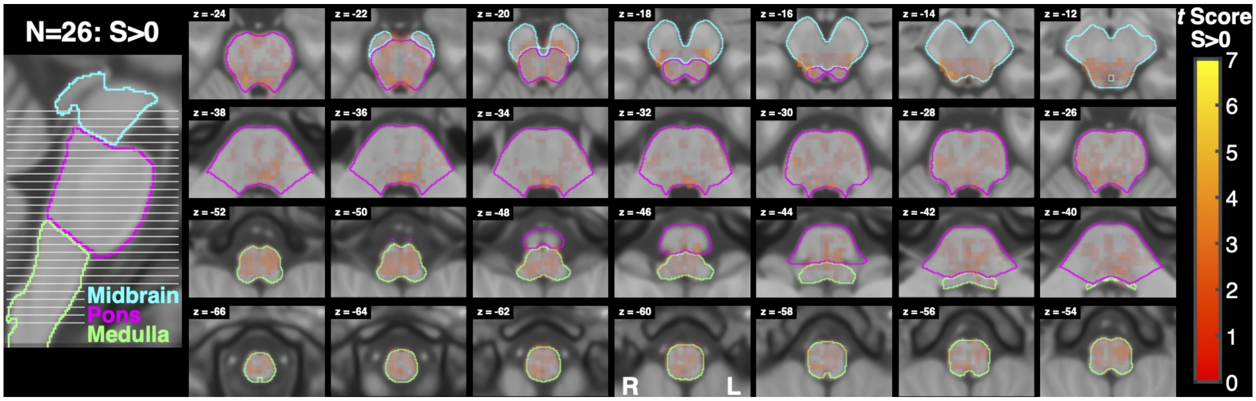


Figure 1: Activation associated with the Slow contrast. A sagittal slice on the left shows the location of each axial slice of the brainstem shown on the right with the highest slice on the top right and the lowest slice on the bottom left. Three sections of the brainstem are outlined with different colors to facilitate orientation. Voxel color maps to $t$ score (no lower limit), while opacity maps to TFCE significance.

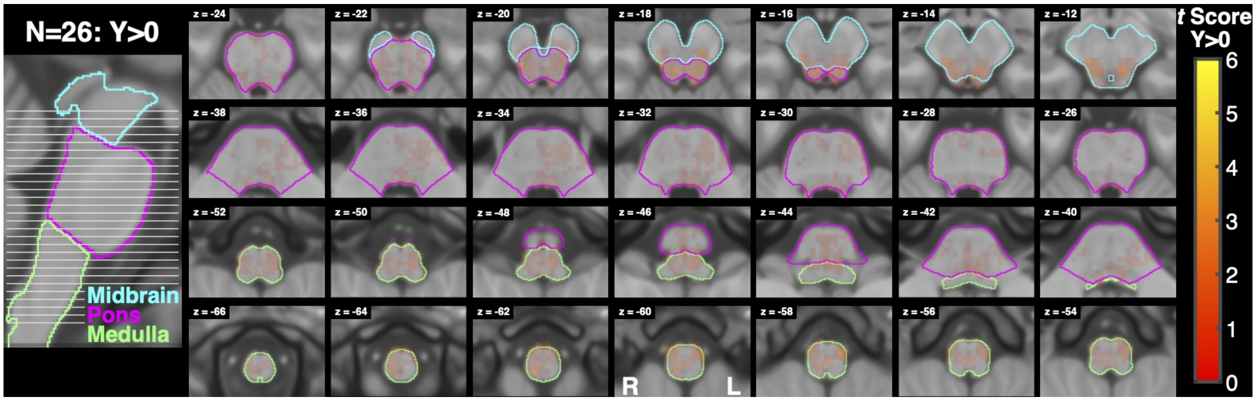


Figure 2: Activation associated with the Yield contrast. A sagittal slice on the left shows the location of each axial slice of the brainstem shown on the right with the highest slice on the top right and the lowest slice on the bottom left. Three sections of the brainstem are outlined with different colors to facilitate orientation. Voxel color maps to $t$ score (no lower limit), while opacity maps to TFCE significance.

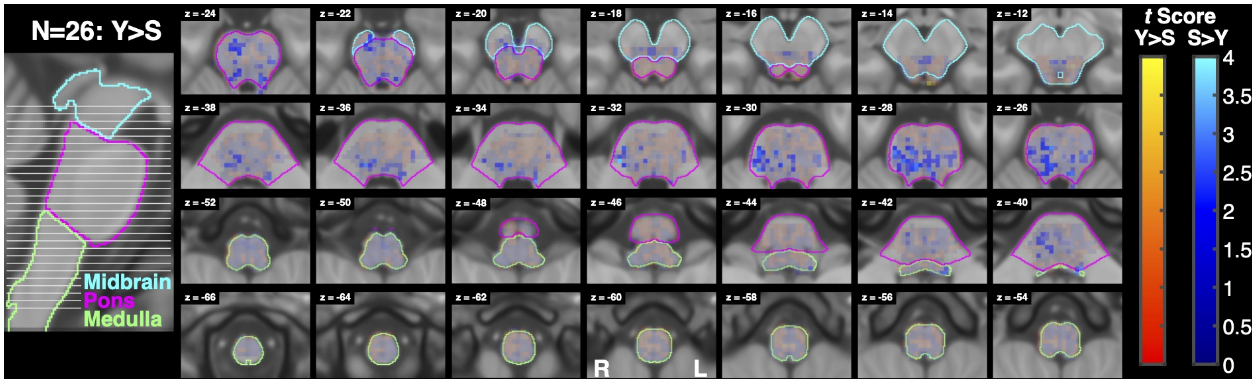


Figure 3: Activation associated with the Yield>Slow contrast. A sagittal slice on the left shows the location of each axial slice of the brainstem shown on the right with the highest slice on the top right and the lowest slice on the bottom left. Three sections of the brainstem are outlined with different colors to facilitate orientation. Voxel color maps to $t$ score (no lower limit), while opacity maps to TFCE significance.

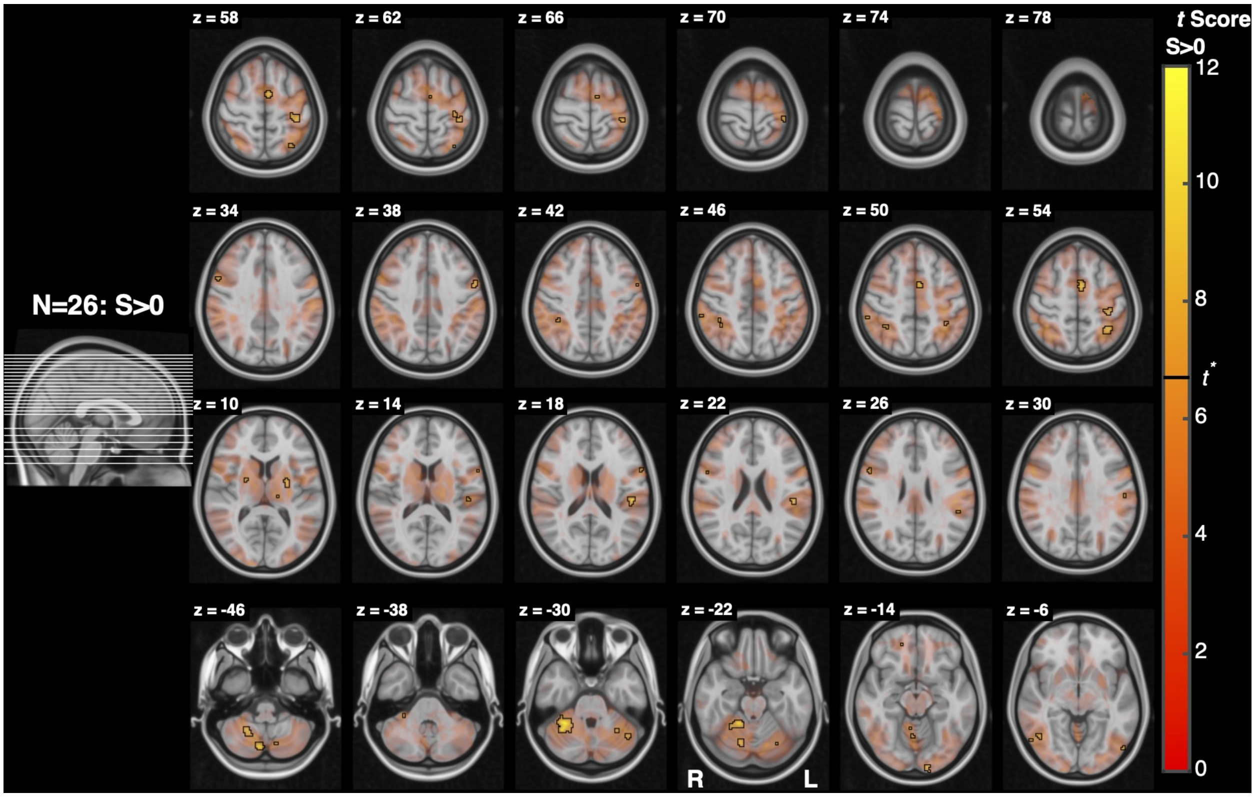


Figure 4: Cortical activation associated with the Slow contrast. A sagittal slice on the left shows the location of each axial slice of the brain shown on the right with the highest slice on the top right and the lowest slice on the bottom left. Functional activation is shown with lower t score voxels in red and higher t score voxels in yellow. Opacity is also modulated by t score and significant voxels are outlined in black.

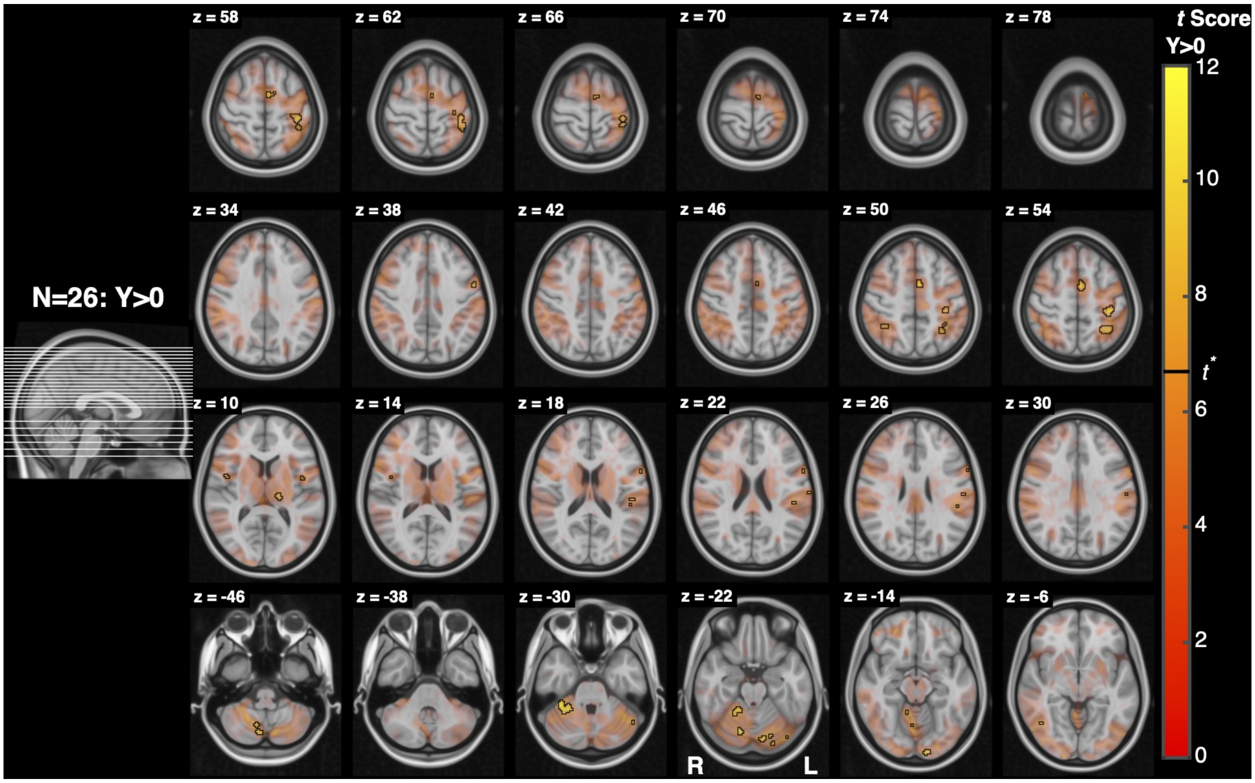


Figure 5: Cortical activation associated with the Yield contrast. A sagittal slice on the left shows the location of each axial slice of the brain shown on the right with the highest slice on the top right and the lowest slice on the bottom left. Functional activation is shown with lower t score voxels in red and higher t score voxels in yellow. Opacity is also modulated by t score and significant voxels are outlined in black.

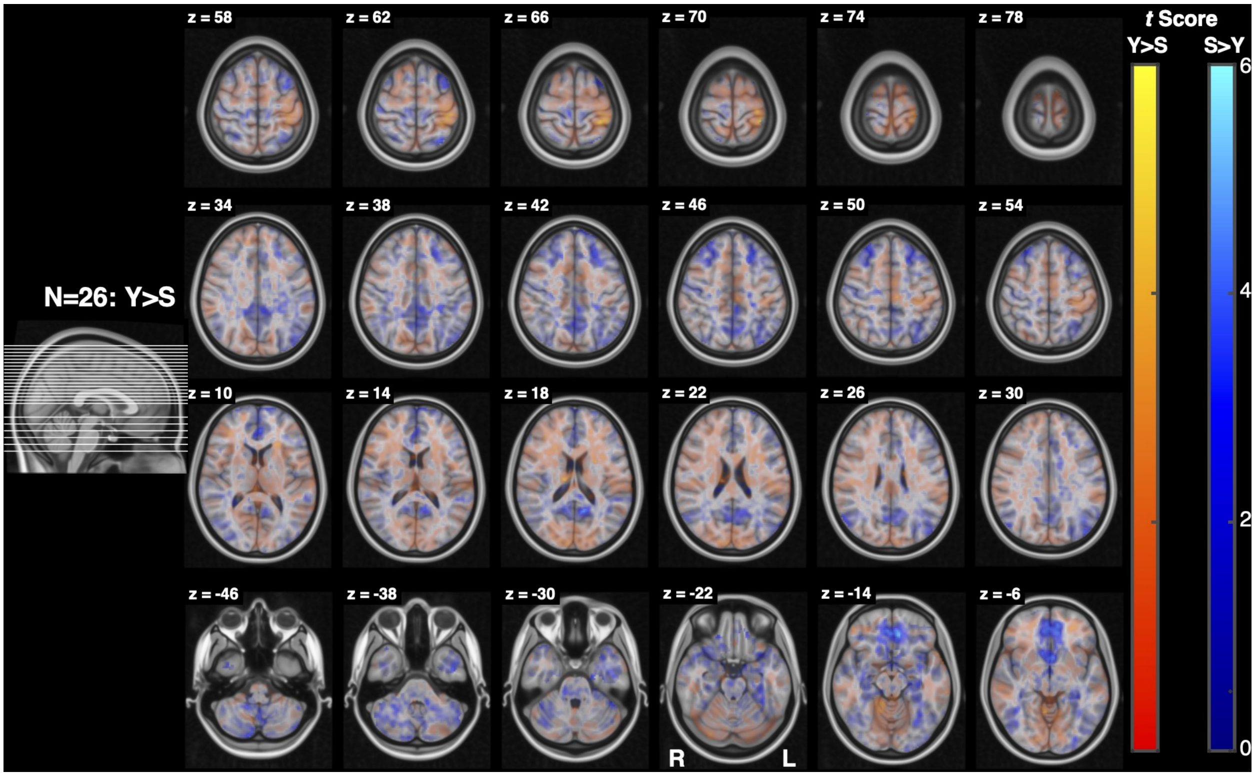


Figure 6: Cortical activation associated with the Yield>Slow contrast. A sagittal slice on the left shows the location of each axial slice of the brain shown on the right with the highest slice on the top right and the lowest slice on the bottom left. Functional activation is shown with lower t score voxels in red and higher t score voxels in yellow. Opacity is also modulated by t score and significant voxels are outlined in black.

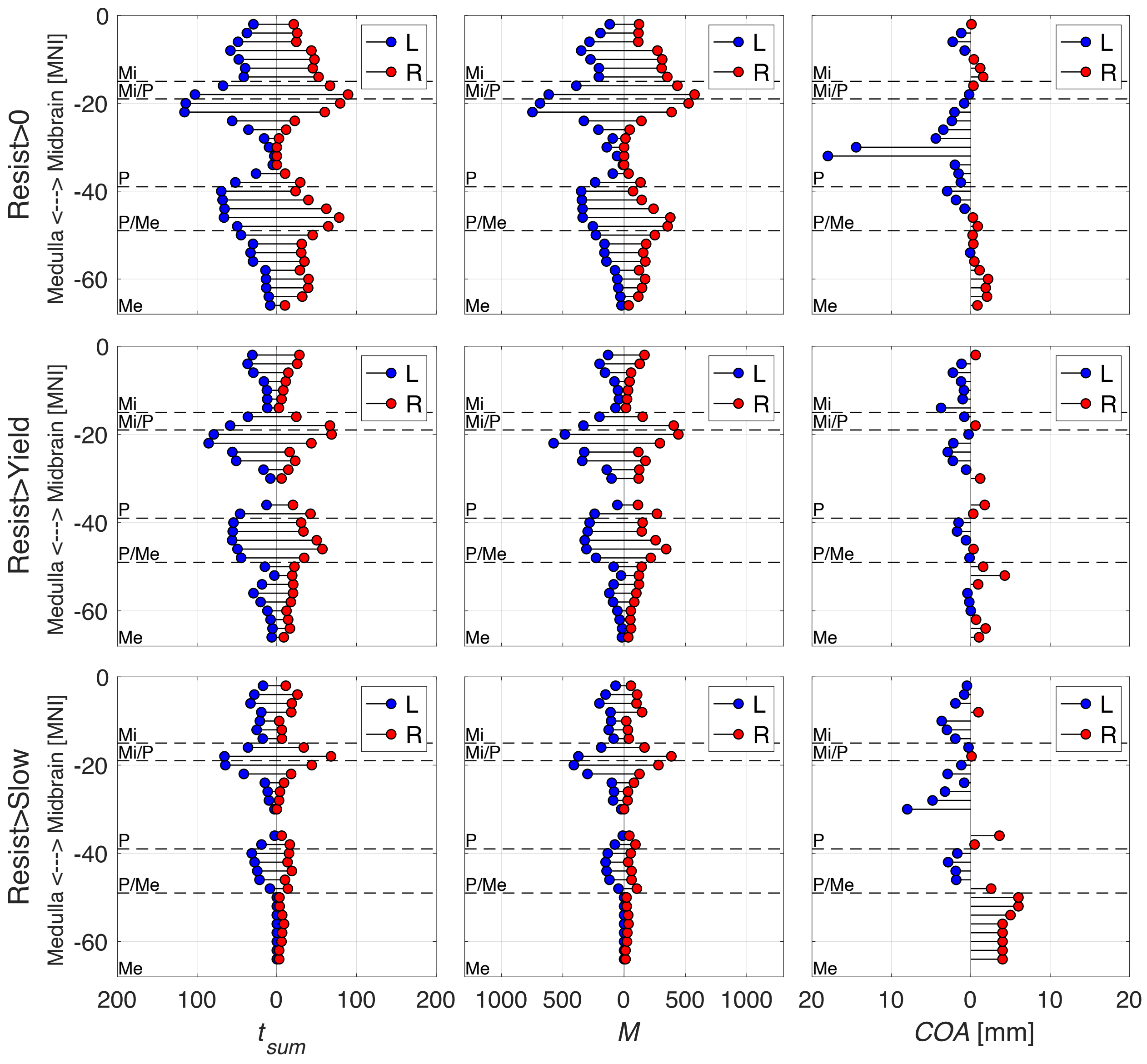


Figure 7: Laterality of significant (TFCE correction) activation in the brainstem. The three methods of quantifying laterality are shown with $t_{sum}$ on the left, $M$ in the middle, and $COA$ on the right with the contrasts Resist on the top, Resist>Yield in the middle, and Resist>Slow on the bottom. Outcome values are on each x axis and axial slice location ($z$ slice in MNI coordinates) are on each y axis. Horizontal dashed lines separate different sections of the brainstem by medulla, pons, midbrain, and their combinations.

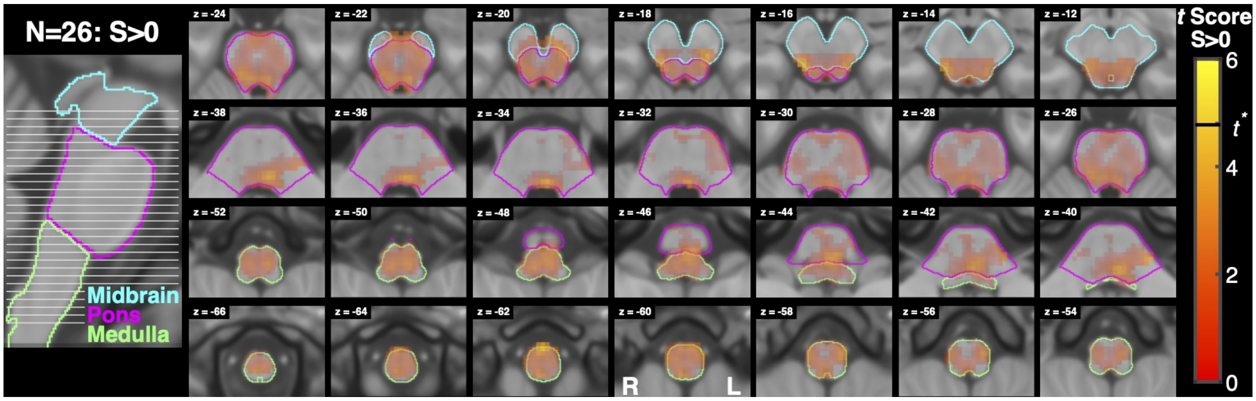


Figure 8: Activation associated with the Slow contrast using the more conventional general linear model without regressors for the rejected MEICA components and 4mm smoothing. A sagittal slice on the left shows the location of each axial slice of the brainstem shown on the right with the highest slice on the top right and the lowest slice on the bottom left. Three sections of the brainstem are outlined with different colors to facilitate orientation. Voxel color and opacity maps to $t$ score (no lower limit), with significant voxels, thresholded using small volume correction, outlined in black. For this analysis $t^* = 4.8$.

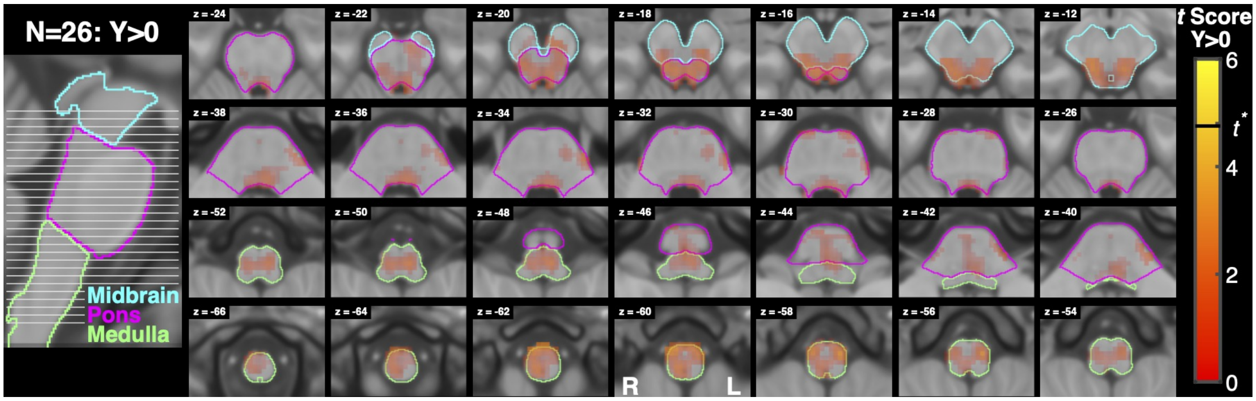


Figure 9: Activation associated with the Yield contrast using the more conventional general linear model without regressors for the rejected MEICA components and 4mm smoothing. A sagittal slice on the left shows the location of each axial slice of the brainstem shown on the right with the highest slice on the top right and the lowest slice on the bottom left. Three sections of the brainstem are outlined with different colors to facilitate orientation. Voxel color and opacity maps to $t$ score (no lower limit), with significant voxels, thresholded using small volume correction, outlined in black. For this analysis $t^* = 4.8$.

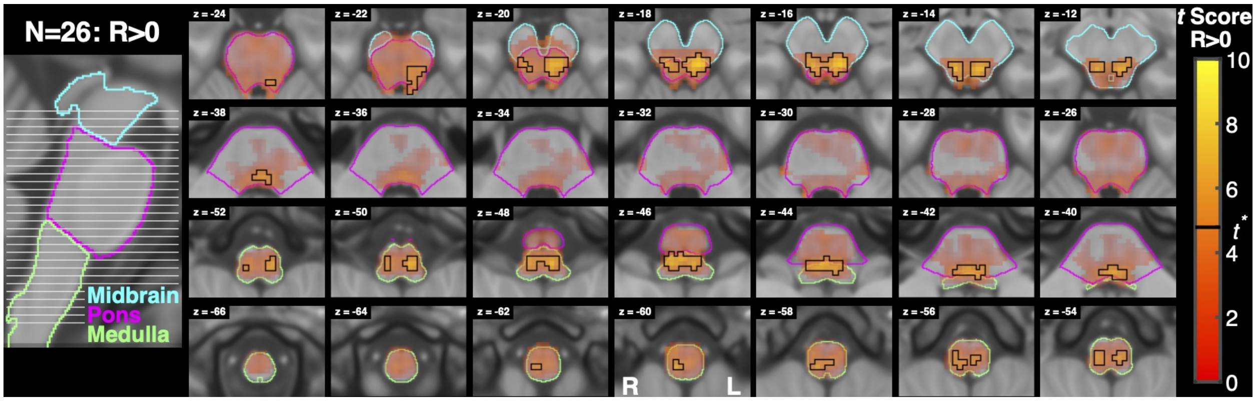


Figure 10: Activation associated with the Resist contrast using the more conventional general linear model without regressors for the rejected MEICA components and 4mm smoothing. A sagittal slice on the left shows the location of each axial slice of the brainstem shown on the right with the highest slice on the top right and the lowest slice on the bottom left. Three sections of the brainstem are outlined with different colors to facilitate orientation. Voxel color and opacity maps to $t$ score (no lower limit), with significant voxels, thresholded using small volume correction, outlined in black. For this analysis $t^* = 4.8$.

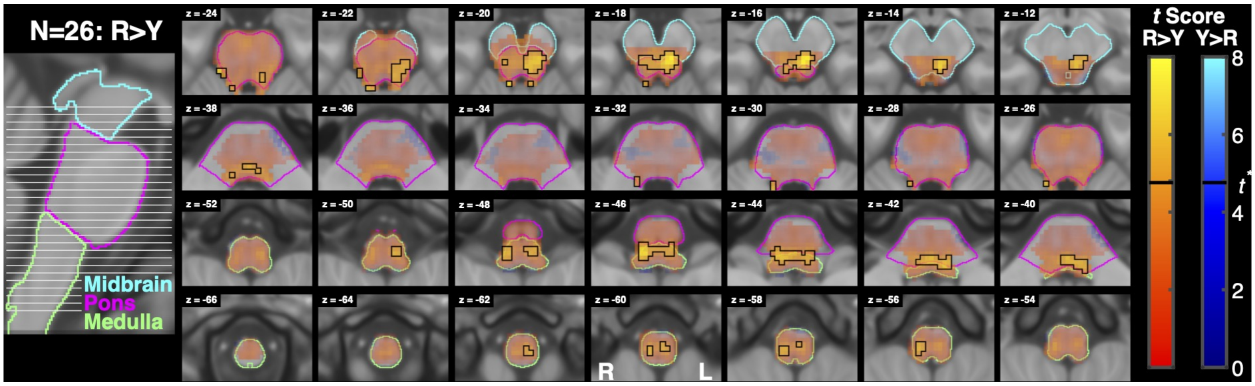


Figure 11: Activation associated with the Resist>Yield contrast using the more conventional general linear model without regressors for the rejected MEICA components and 4mm smoothing. A sagittal slice on the left shows the location of each axial slice of the brainstem shown on the right with the highest slice on the top right and the lowest slice on the bottom left. Three sections of the brainstem are outlined with different colors to facilitate orientation. Voxel color and opacity maps to $t$ score (no lower limit), with significant voxels, thresholded using small volume correction, outlined in black. For this analysis $t^* = 4.8$.

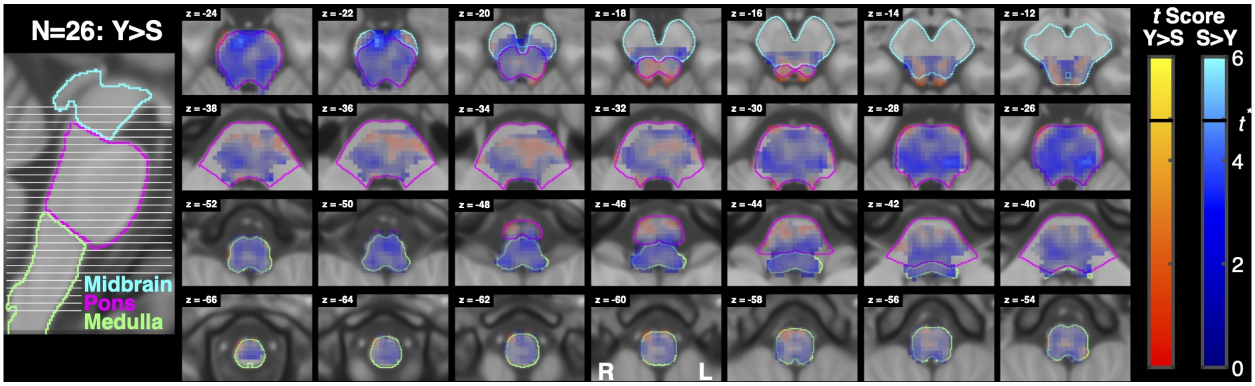


Figure 12: Activation associated with the Yield>Slow contrast using the more conventional general linear model without regressors for the rejected MEICA components and 4mm smoothing. A sagittal slice on the left shows the location of each axial slice of the brainstem shown on the right with the highest slice on the top right and the lowest slice on the bottom left. Three sections of the brainstem are outlined with different colors to facilitate orientation. Voxel color and opacity maps to $t$ score (no lower limit), with significant voxels, thresholded using small volume correction, outlined in black. For this analysis $t^* = 4.8$.

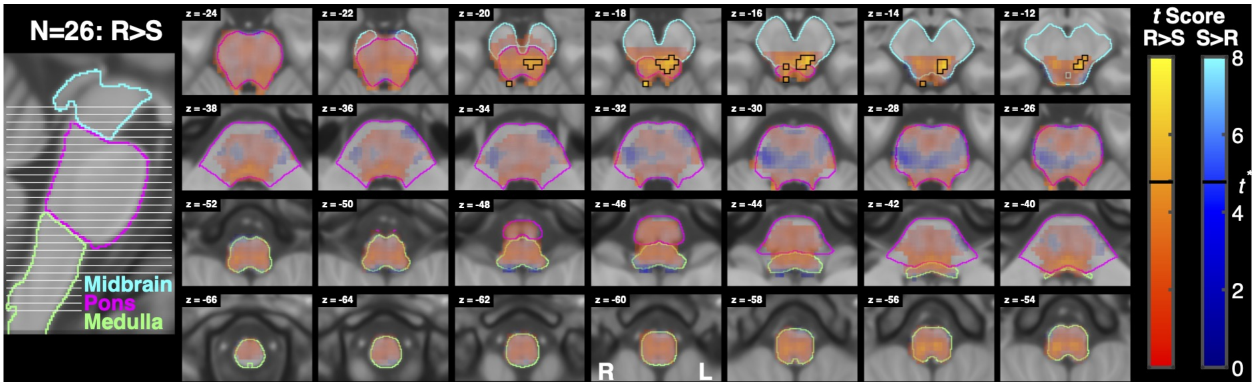


Figure 13: Activation associated with the Resist>Slow contrast using the more conventional general linear model without regressors for the rejected MEICA components and 4mm smoothing. A sagittal slice on the left shows the location of each axial slice of the brainstem shown on the right with the highest slice on the top right and the lowest slice on the bottom left. Three sections of the brainstem are outlined with different colors to facilitate orientation. Voxel color and opacity maps to $t$ score (no lower limit), with significant voxels, thresholded using small volume correction, outlined in black. For this analysis $t^* = 4.8$.
